# DeepFocus: A Transnasal Approach for Optimized Deep Brain Stimulation of Reward Circuit Nodes

**DOI:** 10.1101/2024.10.08.617133

**Authors:** Yuxin Guo, Mats Forssell, Dorian M. Kusyk, Vishal Jain, Isaac Swink, Owen Corcoran, Yuhyun Lee, Chaitanya Goswami, Alexander C. Whiting, Boyle C. Cheng, Pulkit Grover

**Affiliations:** Neuroscience Institute 4400 Fifth Avenue, Carnegie Mellon University, Pittsburgh, PA 15213; Electrical and Computer Engineering, Carnegie Mellon University, 5000 Forbes Ave, Pittsburgh, PA 15213; Allegheny Health Network, 320 E North Ave, Pittsburgh, PA 15212; Biomedical Engineering, Carnegie Mellon University, 5000 Forbes Ave, Pittsburgh, PA 15213

**Keywords:** Neural stimulation, Transnasal electrodes, Optimization, Cadaver studies

## Abstract

**Objective:** Transcranial electrical stimulation (TES) is an effective technique to modulate brain activity and treat diseases. However, TES is primarily used to stimulate superficial brain regions and is unable to reach deeper targets. The spread of injected currents in the head is affected by volume conduction and the additional spreading of currents as they move through head layers with different conductivities, as is discussed in [1]. In this paper, we introduce DeepFocus, a technique aimed at stimulating deep brain structures in the brain’s “reward circuit” (e.g. the orbitofrontal cortex, Brodmann area 25, amygdala, etc.).

**Approach:** To accomplish this, DeepFocus utilizes transnasal electrode placement (under the cribriform plate and within the sphenoid sinus) in addition to electrodes placed on the scalp, and optimizes current injection patterns across these electrodes. To quantify the benefit of DeepFocus, we develop the DeepROAST simulation and optimization platform. DeepROAST simulates the effect of complex skull-base bones’ geometries on the electric fields generated by DeepFocus configurations using realistic head models.

It also uses optimization methods to search for focal and efficient current injection patterns, which we use in our simulation and cadaver studies.

**Main Results:** In simulations, optimized DeepFocus patterns created larger and more focal fields in several regions of interest than scalp-only electrodes. In cadaver studies, DeepFocus patterns created large fields at the medial orbitofrontal cortex (OFC) with magnitudes comparable to stimulation studies, and, in conjunction with established cortical stimulation thresholds, suggest that the field intensity is sufficient to create neural response, e.g. at the OFC.

**Significance:** This minimally invasive stimulation technique can enable more efficient and less risky targeting of deep brain structures to treat multiple neural conditions.

## 1. Introduction

The brain’s “reward circuit” consists of multiple cortical and subcortical nodes (Fig. 1a), including the orbitofrontal cortex (OFC), hippocampus, amygdala, nucleus accumbens, thalamus, etc. This circuitry processes reward and punishment, and is implicated in several conditions relevant to our society, e.g. depression [2, 3], suicidal ideation [4], post-traumatic stress disorder (PTSD) [5, 6, 7], and substance use disorder [8]. Recently, for patients suffering from conditions that are drug-resistant, a promising treatment alternative is emerging: implanted electrodes in nodes of the reward circuit to electrically stimulate these brain areas and successfully treat them. For example, Mayberg et al. used deep-brain stimulation (DBS) to stimulate Brodmann Area 25 (subgenual cingulate area) for patients with drug-resistant depression, and observed remission of depression in four of the six patients recruited [2]. Follow-up studies suggest that the treatment was effective over a long term (i.e., 3-6 years) and did not induce significant adverse effects [9, 10]. Rao et al. [11] stimulated the lateral OFC and observed acute improvement in mood-state in patients with moderate-to-severe depression. Zhang et al. [3] proposed the stimulation of medial OFC as a promising treatment of major depressive disorder (MDD), as conventional antidepressants do not normalize the abnormal activations of medial OFC during MDD. Koek et al. stimulated the basolateral amygdala to treat combat PTSD [12] and Müller et al. stimulated the nucleus accumbens to treat addiction [13].

**Figure 1:**
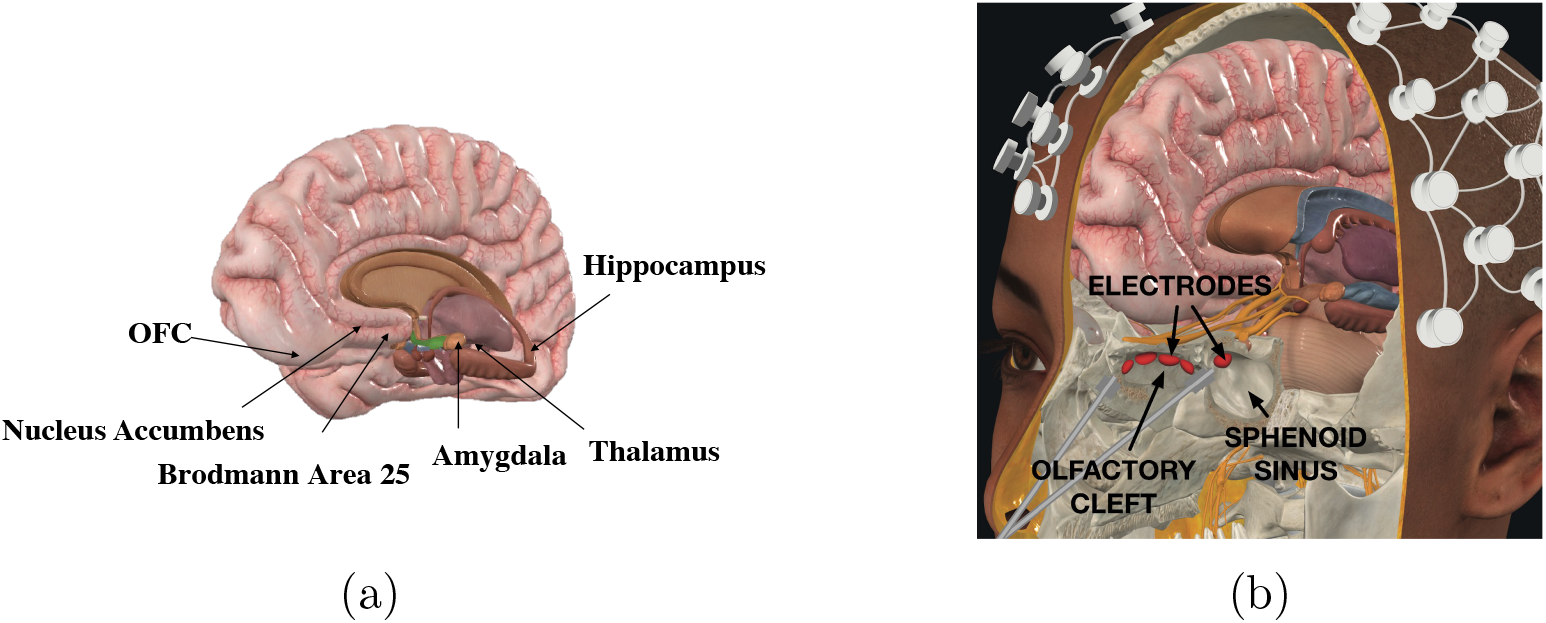
(a) Important nodes in the brain’s reward circuit. These brain regions are involved in reward and punishment processing but are difficult to target with conventional transcranial stimulation due to their inferior locations. (b) DeepFocus places electrodes in nasal cavities, (under the cribriform plate and within the sphenoid sinus), as well as under the scalp, and jointly optimizes injected currents to cause deep stimulation with no shallow stimulation. This figure was created using images courtesy of Complete Anatomy.

However, DBS requires a sophisticated surgical procedure to implant the system (including electrodes, battery, and the associated circuitry) which carries risk of intracranial hemorrhage and infection [14]. Moreover, the stimulation target region cannot be changed once the electrodes are implanted inside the brain. Non-invasive techniques such as transcranial magnetic stimulation (TMS), transcranial focused ultrasound stimulation (tFUS), and transcranial electrical stimulation (TES) have lower risk and are steerable, but are also limited in their depth, resolution, and effect [1, 15, 16]. TMS and pulsed TES (at high currents and short pulse widths [17]) can directly stimulate neurons, resulting, for example, in motor evoked potentials when targeting the motor cortex. However, when used to target deep brain structures, they stimulate superficial structures more powerfully than deep targets and require higher currents, leading to potential side-effects, including high scalp pain [18, 19]. A recent discovery of temporal interference (TI) stimulation is interesting in this context, and is discussed in detail in Section 5.2. The mechanisms of tFUS are not well understood, and its effects are thought to cause potentiation or modulation of firing rate rather than direct stimulation [20].

In this paper, we introduce the DeepFocus technique, which places electrodes transnasally (see Fig. 1b) in the olfactory cleft and within the sphenoid sinus to perform transnasal electrical stimulation (TnES), in addition to electrodes on the scalp for transcranial electrical stimulation (TES). Further, DeepFocus optimizes current injection patterns for efficient and focal targeting of reward circuit nodes. In placing electrodes inside the olfactory cleft (under the cribriform plate) and within the sphenoid sinus, we aim to harness the physical proximity to target deep brain structures, as well as the high-conductive pathways offered by thin bones and foramina in the cribriform plate.

To perform this optimization and quantify the benefit of DeepFocus over traditional scalp-only current injection patterns, we develop the DeepROAST platform (Section 2). Our platform builds on a recent advance – ROAST [21, 22] – which is an integrated platform for image segmentation, meshing, and solving for fully-automated field simulations resulting from TES. DeepFocus also utilizes optimization techniques that have been developed for focal transcranial stimulation [23, 24, 25, 26, 27, 28]. These techniques search for optimized electrode placement on the scalp that induce desired fields inside the brain.

To quantify the benefits of DeepFocus, we compare the electric fields generated by using DeepFocus and traditional scalp-only electrodes stimulation. We quantify the gain of DeepFocus in intensity and focality in both simulations (Section 3) and cadaver studies (Section 4). We conclude that the DeepFocus technique creates larger and more focal fields at deep brain targets than traditional scalp electrode configurations. Moreover, the stimulation is steerable, i.e., the clinicians can choose and change the target after electrode placement, which is an advantage over implantable deep brain stimulation.

### Feasibility of transnasal electrode placement

#### Olfactory cleft

The olfactory cleft is close to the ventral side of the brain and can be accessed through the nose with endoscopic guidance. This space is directly below the cribriform plate, which is a thin and complicated bone structure with foramina (i.e., holes) that allow blood vessels and olfactory nerves to pass through. Directly above the cribriform plate is the olfactory bulb, which is situated below the medial orbitofrontal cortex of the brain. Previous work has demonstrated safe insertion of electrodes transnasally in the olfactory cleft, and delivery of currents using these electrodes. Authors in [29, 30, 31, 32, 33] inserted stimulation electrodes inside the olfactory cleft to stimulate the olfactory bulb for smell sensations. Of particular interest is the work of Holbrook et al. [29], who placed electrodes directly under the cribriform plate, just like our studies, and injected current up to 15 mA. Weiss et al. [33] also stimulated the olfactory bulb using transnasal electrodes, and additionally recorded brain activities with fMRI. Although they failed to induce odor perception, they found that transnasal olfactory bulb stimulation significantly decorrelated left from right primary olfactory cortex and hippocampal connectivity, which implies that stimulation of the olfactory bulb can *indirectly* alter deep brain activity. Note that for all of these studies, although transnasal electrodes were used for stimulation, the objective was the stimulation of the olfactory bulb and not (direct) deep-brain stimulation.

#### Sphenoid sinus

The sphenoid sinus is an air-filled structure proximal to the pituitary gland and the cranial nerves. The sphenoid sinus is accessible with minimally-invasive procedures. Although no prior work has attempted to stimulate neurons through the sphenoid sinus, FDA-approved sinus implants that release steroids are commonly used to enhance sinus surgery outcomes [34].

A closely related question is: how difficult is this procedure? ENT surgeons routinely access the olfactory space, e.g. for sinus surgeries. Local anesthetics such as tetracaine are frequently used to reduce discomfort. Specific to electrode placement, recently, Menzel et al. [35] investigated the surgical procedure of transnasal electrode placement directly under the cribriform plate in fresh human cadavers. They concluded that endoscopic-guided transnasal positioning is relatively easy and quick (require 15 minutes for experienced ENT surgeons, after preparing the olfactory mucosa) to perform. Placement in the sphenoid sinus is, however, harder. The access to the sinus is through the sphenoid ostium, an opening into the sinus that is accessible transnasally, but could require soft-tissue manipulation to insert an electrode. This is precisely what might make DeepFocus minimally-invasive, instead of non-invasive.

## 2. Materials and Methods

### 2.1. Head Model

The simulation was conducted on a head model with realistic head geometry and tissue conductivity. The New York head model [36], with 0.5mm resolution was used to construct the forward matrix, which relates the applied current at scalp electrodes to the electric field in brain voxels. Since the transnasal electrodes were placed inside the olfactory cleft, realistic modeling of the ethmoid bone anatomy is necessary. However, anatomical details are typically not preserved after tissue auto-segmentation, so we manually modified the New York head tissue masks, specifically the cribriform plate, with reference to existing literature that informs the area and number of the cribriform plate foramina [37, 38]. The size of the foramina was assumed to be 1 mm^2^, with 5 on the left (total area = 5 mm^2^) and 3 on the right (total area = 3 mm^2^) segment of the cribriform plate. The conductivities of tissues are 0.126, 0.276, 1.65, 0.01, 0.465, 2.5 *×* 10^−14^ S/m for white matter, gray matter, CSF, bone, skin, and air respectively [21]. In addition, since olfactory nerves pass through the cribriform foramina, the corresponding voxels were assigned the conductivity of nerves (0.39 S/m) [39]. The tissue segmentation is shown in Figure 2.

**Figure 2:**
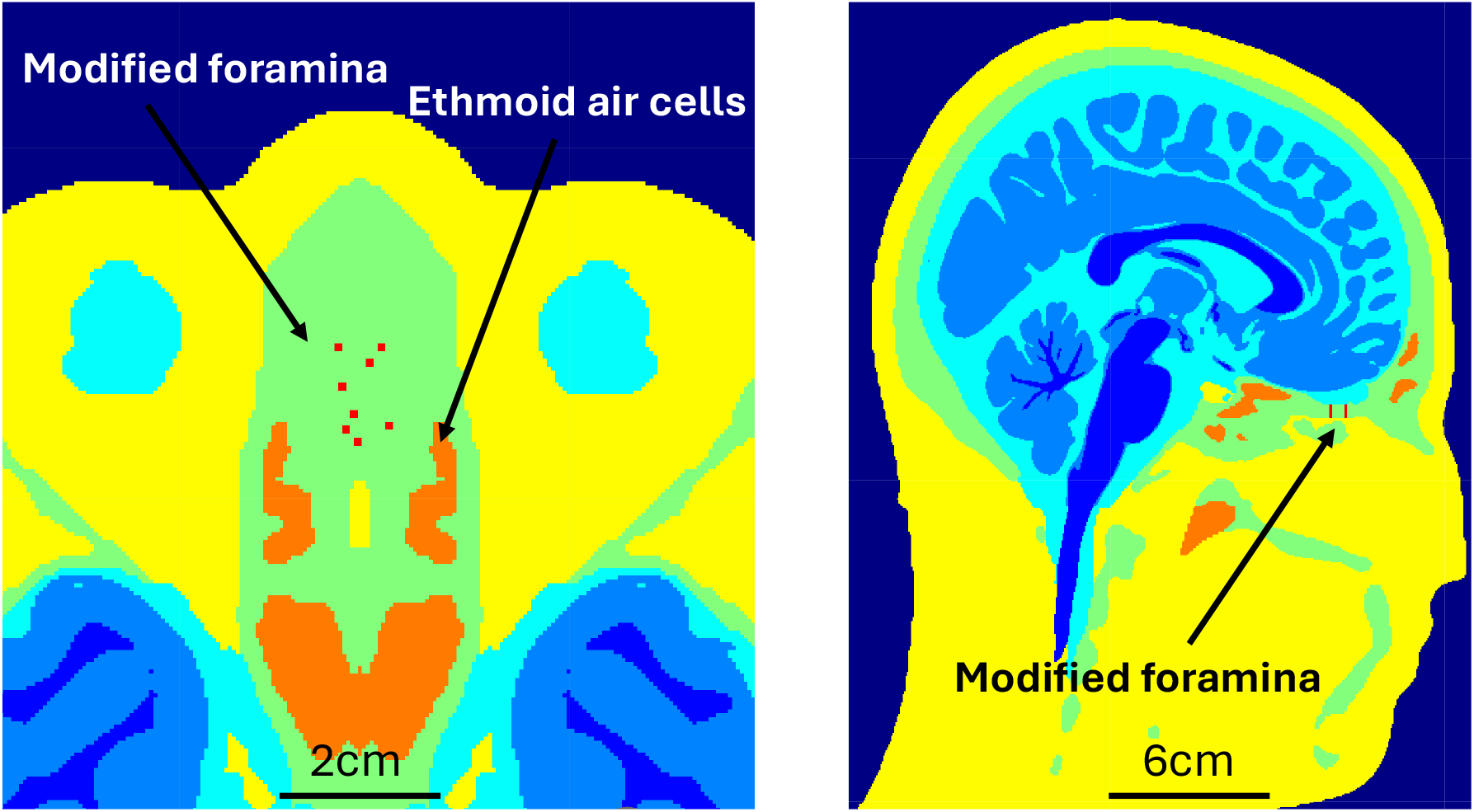
Horizonal and sagittal views of segmented head and brain tissues after addition of foramina on the cribriform plate. The air-filled ethmoid labyrinth, sphenoid sinus, and frontal sinus are also shown in orange. The colors indicate {dark blue: white mater, blue: gray matter, cyan: CSF, green: bone, yellow: skin, orange: air, red: nerve} respectively.

### 2.2. Placement of Electrodes in Simulation

Thirty-two electrodes were placed on the scalp according to the “10-20” system [40]. These scalp electrodes are modeled as disk electrodes with a radius of 4 mm. In addition, 5 electrodes were placed inside each half of the sphenoid sinus with a spacing of at least 5 mm, adjusting as required to the sphenoid sinus anatomy, for placement in contact with the sinus wall. 4 electrodes were placed beneath the cribriform plate bilaterally in each hemisphere (8 total), separated by 5 mm along the anterior-posterior direction. The electrodes on both sides of the olfactory cleft were separated by the perpendicular plate of the ethmoid bone, with a distance of 1 cm. The transnasal electrodes were modeled as disk electrodes with a radius of 2 mm.

### 2.3. Simulation Pipeline

The forward simulation was completed with the ROAST platform with adaptations described above that allowed customized placement of transnasal electrodes and more realistic conductivity of skull-base bones. The automatic pipeline includes tissue segmentation from MRI scans, conductivity value assignment, electrode placement, 3D meshing, and solving with the finite element method (FEM).

FEM simulations were used to solve the Laplace equation, where the electric field along each of *x, y*, and *z* direction at brain voxels 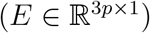 and currents applied by electrodes (*s* ∈ ℝ^*n×*1^) are related by:

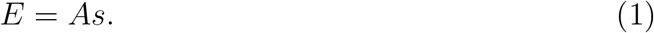

Here, *n* is the number of electrodes and *p* is the number of brain voxels. Since the total inward current must equal the total outward current, the entries of *s* must sum up to 0. 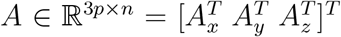 is the forward matrix derived from concatenating the results of the *n* forward simulations column-wise [25], with *A*_*x*_, *A*_*y*_, and *A*_*z*_ indicating the forward models along *x, y*, and *z* directions. In our simulations, we use 18 transnasal electrodes (8 in the olfactory cleft and 10 in the sphenoid sinus, split equally in the two hemispheres) and 32 scalp electrodes, for a total of *n* = 50 transnasal and scalp electrodes. Because our targets are within the brain, the forward matrix only contains voxels in the brain (segmented as either gray matter or white matter), which leads to a total number of voxels *p* = 13, 246, 011.

### 2.4. Optimization for Maximum Intensity at the Target

For simplicity, we assume that the region of interest (ROI) is a single voxel in the target region. To target the ROI with a large electric field along the desired direction (regardless of fields in other regions), the following optimization framework is used [25, 24, 41]:

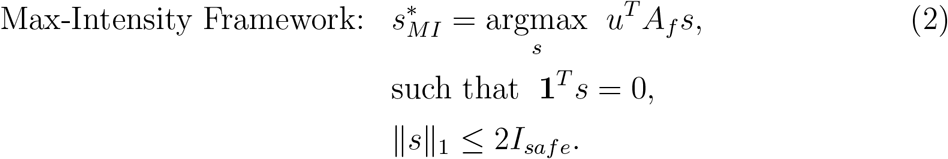

The matrix *A*_*f*_ ∈ ℝ^3*×n*^ are the rows of *A* corresponding to the ROI voxel only. When multiplied by *s*, it yields the electric field intensity in *x, y*, and *z* direction at the ROI. The unit vector *u* is the desired direction of the stimulation electric field, and taking dot product yields the electric field intensity along *u*. In our tests, the direction vector *u* is randomly sampled. The constraints make sure that Kirchhoff’s current law is satisfied (i.e., the currents at the anode and cathode electrodes sum up to 0) and the total injected current is bounded by a safety limit (which yields an *L*1 constraint).

### 2.5. Optimization for Focality of Stimulation

In order to create focal fields at the target, the magnitude of fields at non-target regions should be kept small. The following convex optimization framework is used to encourage focality [24, 28].

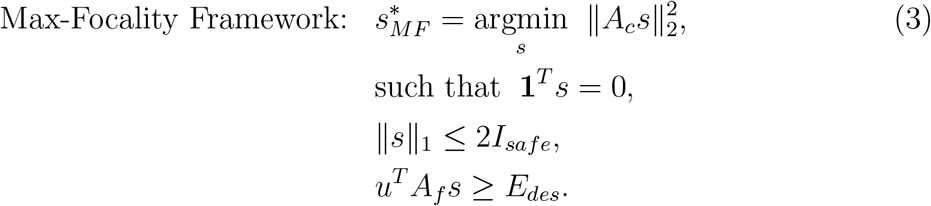

In the objective, *A*_*c*_ ∈ ℝ^*m×n*^ is the forward matrix corresponding to the cancel region, where the field magnitude within this region should be minimized. Minimizing the L2 norm of *A*_*c*_*s* minimizes the (non-directional) magnitude of electric fields at the cancel region. The additional equality constraint specifies that the directional intensity at ROI voxels reaches a desired value *E*_*des*_ (e.g. the neural threshold for stimulation). While in Section 2.4, the maximum intensity along the desired direction is optimized in the objective, here it is manually input into the constraint, thus should be utilized if *E*_*des*_ is known. *E*_*des*_ determines the operating point on the intensity-focality trade-off: small *E*_*des*_ results in focal fields that are lower in intensity, and large *E*_*des*_ induces high-intensity but diffused fields [1, 24, 25]. Note that, if *E*_*des*_ is too large, specifically, larger than the objective value of Max-Intensity Framework 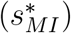,the optimization problem becomes infeasible due to the total current constraint.

In practice, one may be interested in constraining the current density instead of the total injected current. In that case, the L1 constraints in both optimization frameworks can be replaced by 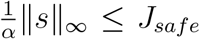,where *J*_*safe*_ is the safety limit for current density, and *α* is the area of the electrode. This constrains the maximum current density by a safety limit. DeepROAST platform supports the options of both total current and current density constraints for the optimization frameworks.

Since the objective is quadratic and the number of voxels is large, the optimization problem is computationally expensive. For computational efficiency, we performed singular value decomposition of the matrix *A*_*c*_ for optimization (i.e., *A*_*c*_ = *U* Σ*V*), as was done in [24].

### 2.6. Quantitative Metrics for Intensity and Focality

In Section 3, we quantify and compare the performance of DeepFocus and scalp-only configurations in terms of intensity and focality. We target some of the reward circuit nodes and compare the results achieved by DeepFocus and scalp-only patterns. DeepFocus has 50 electrodes in its search space, while scalp-only configuration has 32 scalp-electrodes. Stimulation performance is assessed based on metrics of “Intensity Gain” and “Focality”. The Intensity Gain is the ratio of the electric-field intensity attained by DeepFocus and scalp-only, i.e., *E*_*DeepF ocus*_*/E*_*scalp only*_ at ROI along the desired direction.

Focality is quantified using three metrics. First, we define *PV ER*, the percentage of voxels, inside the brain and within the cancel region, with electric-field magnitude exceeding that of the ROI voxel. This metric is analogous to the commonly used full-width-half-max (FWHM), but is more suited to our problem, since the overall field maximum might not be attained in the ROI (e.g. when the ROI is not adjacent to any electrode). Second, similar to [24], we compute *I*(*k*), the integral of the electric field magnitude included within the *k*-nearest brain voxels of the ROI, and report *I*_*frac*_(*k*) = *I*(*k*)*/I*(*p*), the ratio of the integral *I*(*k*) to the integral of the field in the entire brain (note that field outside brain voxels is not included in this integral). Finally, we also compute the *K*-value (*K*_50_) from the second metric, where *K*_50_ is the threshold value of *k/p* for which *I*_*frac*_(*k*) = 0.5, i.e., the fraction of voxels nearest to the ROI such that 50% of the total field is reached. We report *K*_50_ as a percentage.

For consistency, the total injected current (i.e., *I*_*safe*_) is fixed to 1 mA in all simulations.

### 2.7. Monte Carlo Simulations

The cribriform plate of the ethmoid bone has a lot of variation across individuals, including the size and the number of foramina [37]. In Section 3.4, we perform Monte Carlo simulations to understand how the variations in the cribriform plate anatomy can affect our results. We modified the NY Head tissue mask for the cribriform plate foramina, where the total area of foramina on each side of the cribriform plate is drawn uniformly in the interval [3, 9] mm^2^ [37]. The cribriform plate was segmented, and 10 sets of foramina were generated at random locations on the cribriform plate. We include the visualization of the Monte Carlo samples in the supplementary material Section 3. Due to their location and proximity to olfactory cleft electrodes, electric fields in the brain region close to the cribriform plate, namely, the medial anterior OFC, is expected to be most sensitive to changes in cribriform plate anatomy. Therefore, in the Monte Carlo simulations, we used the DeepFocus pattern with an anode at the olfactory cleft electrode and the cathode at Fz, targeting medial anterior OFC.

### 2.8. Cadaver Experiment Setup

To test the efficiency of DeepFocus, we performed cadaver studies with specimens obtained from *Anatomy Gifts Registry*. The cadaver tissue specimen used in this study was obtained from a 54 year-old-male donor. Two experiments were performed using the same specimen. In the first experiment to target the medial OFC, a neurosurgeon (D.K.) inserted four stereoencephalography (sEEG) probes (Ad Tech Medical Instrument Corp.), roughly 1 cm apart bilaterally in the medial OFC, reaching just above the cribriform plate, for field recordings. Each sEEG probe contains 10-12 cylindrical electrodes, at a 5 mm pitch. Transnasal electrodes were inserted with endoscopic guidance in the left olfactory cleft above the superior turbinate, directly below the cribriform plate, as shown in Fig. 3a. The placement of electrodes was verified by 3d x-ray imaging (O-arm, Medtronic). Gold-cup electrodes were placed on the scalp at locations FP2 and P7 according to the EEG 10-20 system.

**Figure 3:**
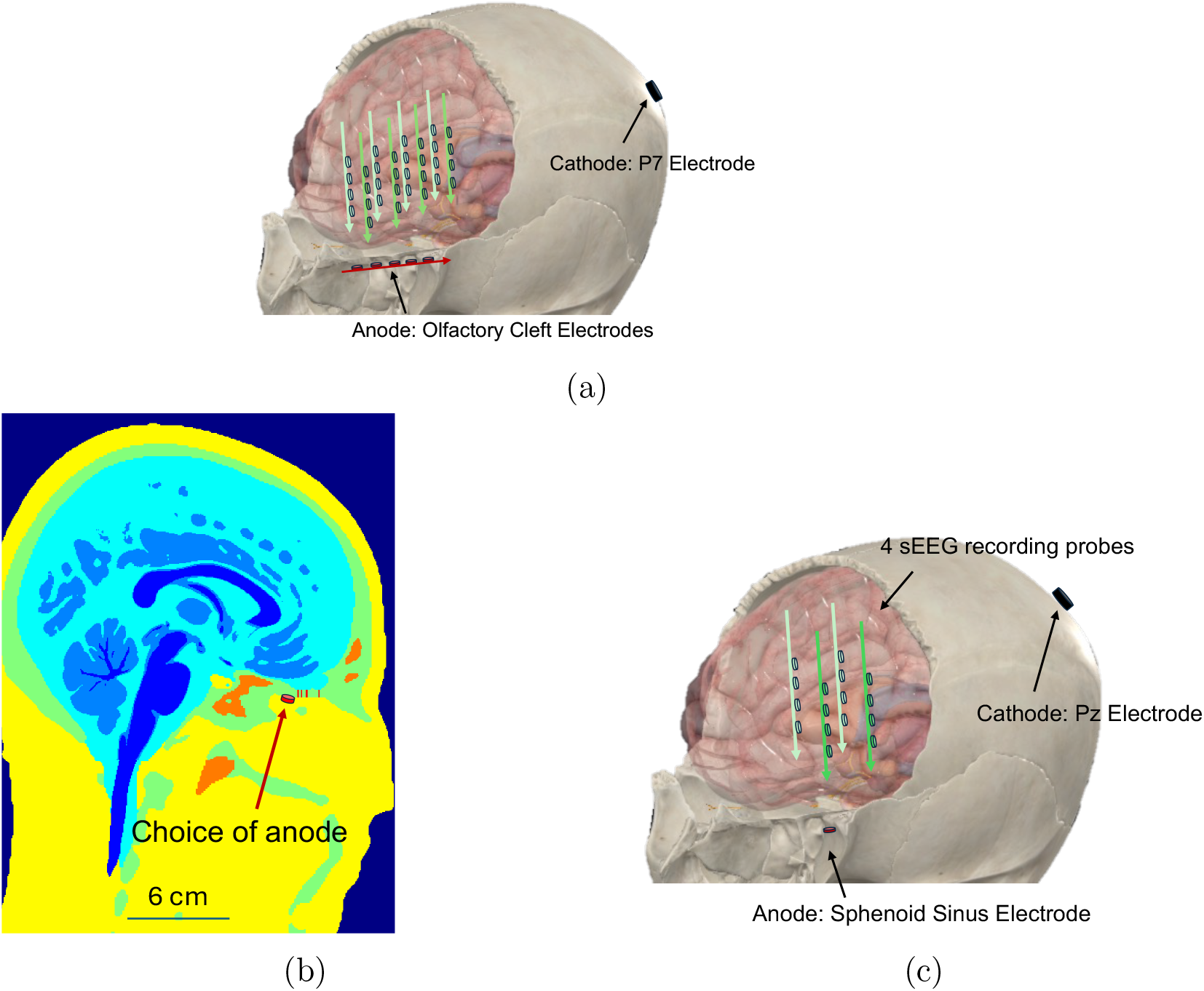
Placement of electrodes in the cadaver specimen demonstrated in the New ^1^ York head and Complete Anatomy models. Images are not reflective of the cadaver-specimen-specific anatomy. (a) Eight recording stereoelectroencephalography (sEEG) probes were inserted bilaterally (i.e., four in each hemisphere) to reach the medial OFC from above. The stimulation sEEG electrodes were inserted through the left nostril below the cribriform plate using endoscopic guidance. The cathode P7 was placed according to the 10-20 system. (b) The anode location inside the olfactory cleft was chosen such that it was in close proximity of the cribriform plate. (c) The anode was placed directly under the superior wall of the sphenoid sinus (shown in red), and the cathode was placed at Pz (shown in blue). sEEG electrodes are shown in green. (a) and (c) were created using images courtesy of Complete Anatomy, (b) is from the New York head model.

In the second experiment, designed to target the ventral diencephalon, four sEEG probes were repositioned to reach the ventral diencephalon in the left hemisphere (as shown in Fig. 3c). We inserted an electrode transnasally into the left sphenoid sinus, in contact with the superior wall of the sphenoid sinus. The cathode was a gold cup electrode placed on the scalp at Pz. All procedures were approved by the Department of Anatomic Pathology and Clinical Pathology at Allegheny Health Network (Anatomic Materials Protocol No.: AM-2023-2).

In each experiment, stimulation pulses were applied to selected electrode pairs (nasal electrode and/or scalp electrode), while the resulting voltage at the brain electrodes was recorded. Biphasic, 10 ms/phase pulses of 10 mA amplitude were injected using a commercial neurostimulator (DS5 stimulator, Digitimer Ltd.). Pulses were repeated at 10 Hz, and the resulting fields averaged over 300-600 pulses. Due to limitations of the recording instrumentation, we could only record the voltage from 16 channels simultaneously. Therefore we only measured the voltage from the four inferior electrodes on each probe. In the experiment to target the medial OFC, two recordings were performed and concatenated to measure potentials from the inferior portions of all eight probes.

The recordings were analyzed using custom scripts in MATLAB. The amplitude of the voltage pulse resulting from the stimulation pulse was calculated for each electrode. The exact location of each electrode was manually measured from the 3D x-ray images. The evoked potential field was then reconstructed using linear interpolation followed by a 3D Gaussian smoothing filter with standard deviation of 5 mm. The electric field inside the brain was estimated as the gradient of the measured potentials. For consistency with the simulation results we scaled the fields in Fig. 10 by a factor of 0.1 to report the field created by a 1 mA stimulation current.

In addition, we conducted a comparative analysis between the cadaver and simulation results. The brain volume within which the electrodes were inserted during the cadaver experiments was mapped onto the New York head model by visually inspecting the CT scans. To quantify intensity, we calculated the maximum electric field magnitudes from both the simulation and cadaver results, constrained within the respective volumes. To quantify focality, we calculated the focality metric *I*_*frac*_ for both the simulation results and the cadaver experiment recordings. For the sphenoid sinus stimulation, *I*_*frac*_ was calculated with the electric field magnitude along the inferior-superior direction. This approach was chosen because only a 2 *×* 2 grid of recording probes was inserted, leading to coarser measurements in the medial-lateral and anterior-posterior directions.

## 3. Simulation Results

### 3.1. Orbitofrontal Cortex

Targets of stimulation were sampled from the anterior, posterior, medial, and lateral OFC at random locations. For each region, 10 randomly sampled target ROIs (one voxel each) were utilized in our estimates, and the statistics of their intensity gains are shown in Section 3.3. The ROI of each sample is a single voxel within the region, and the optimal direction for each sample is a random three-dimensional unit vector.

#### Intensity Gain

To quantify the Intensity Gain, we utilized the Max-Intensity Framework as described in Section 2.4 for optimization. Because DeepFocus places electrodes in close proximity to the medial-posterior OFC, it can be targeted with the largest field intensity. In contrast, with scalp-only configurations, medial-posterior OFC has very small field intensities. The maximum intensity gain is 127.6 at a medial-posterior ROI. To target that ROI voxel, a combination of electrodes from sinuses on both sides are used, and the resulting electric field is shown in Figure 4. In scalp-only configurations, optimization with Max-Intensity Framework utilizes the forehead electrodes F7 and F8 to target the ROI. Despite no explicit constraints of focality in the optimization framework, DeepFocus creates more focal field at the OFC, and scalp-only configuration activates superficial regions outside the target mostly near the electrodes, as shown in supplementary material Fig.13.

**Figure 4:**
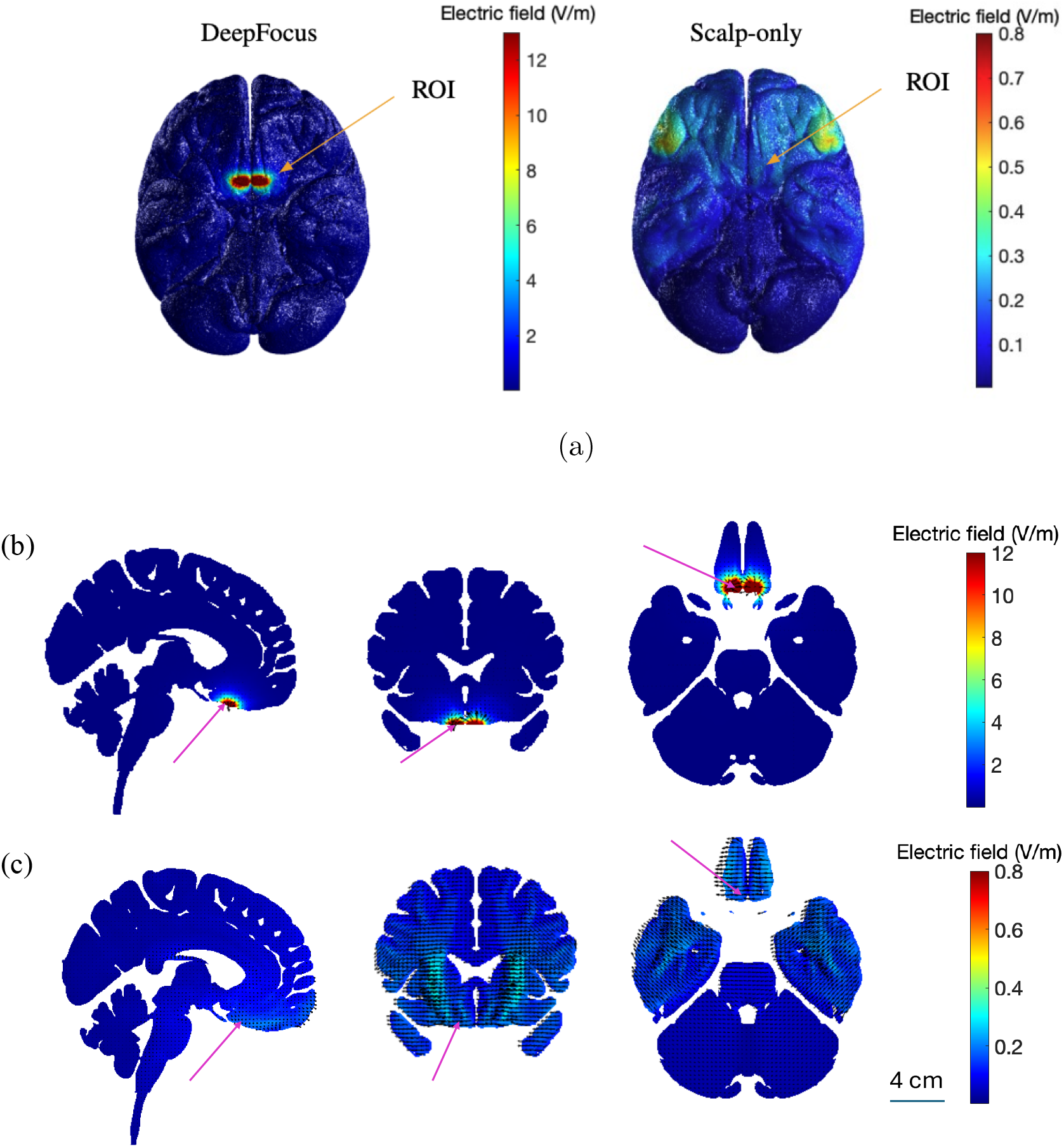
Intensity gain at an OFC target. Due to the *L*1 constraint in the Max-Intensity framework that encourages sparsity, the optimizing electrode configurations often utilize two electrodes (i.e. one anode and one cathode). (a) The magnitude of the electric field created by DeepFocus Max-Intensity framework and scalp-only patterns. DeepFocus utilizes two electrodes, both in the sphenoid sinus (one in each side of the sinus), and the scalp-only configuration utilizes F7 and F8. The total injected current is 1 mA in both cases. (b) Slice view of the electric field generated by the DeepFocus pattern. Slice view of the electric field generated by the scalp-only pattern. The DeepFocus pattern creates a much larger and more focal field at the ROI, whereas the scalp-only pattern produces a larger field in more shallow regions, with a smaller intensity field at the ROI.

#### Focality

Next we investigate focality-constrained optimization for the 10 randomly selected OFC targets. We employed the optimization framework in Section 2.5. We keep our ROI to be a single voxel, and the cancel region (i.e., voxels with minimized field) to be the voxels excluding the ROI and its *k*-nearest voxels. For the reported results, we specified *k* = 1000, which is equivalent to a volume of 125 mm^3^. Fig. 5a shows the *PV ER* for both scalp-only and DeepFocus configurations as the desired intensity increases along the x-axis. The electric fields created by scalp-only electrodes are more diffused than those by transnasal configurations. Additionally, scalp-only optimization problems are unable to reach the desired intensity at ROI with a much lower target intensity value, given the total current constraint. Fig. 5b shows the percentage of fields included in an increasing volume around the ROI at OFC (i.e. *I*_*frac*_(*k*)). The desired intensity at the target is set to a nominal 0.1V/m, corresponding to the largest intensity that is feasible using scalp-only configurations. Since the electric field scales linearly with injected current, if the current in all electrodes is scaled by a constant (same constant across all electrodes), the electric field is scaled by the same factor. We observe that the *PV ER* is much larger with scalp-only configurations, indicating more diffused fields, as shown in Fig. 5a. In Fig. 5b, the curve corresponding to DeepFocus has a much larger slope at smaller intensities and plateaus faster than scalp-only configurations, indicating that DeepFocus fields concentrate much more around the ROI, and thus have better focality. This is also illustrated in Fig. 5c, where the *K*_50_ value is lower for DeepFocus than for scalp-only configurations, which indicates that 50% of the total field magnitude concentrates in a more localized brain volume. The electric field created by both scalp-only and DeepFocus patterns with an intensity of 0.1 V/m is included in the supplementary material (Fig. 11 and Fig. 12).

**Figure 5:**
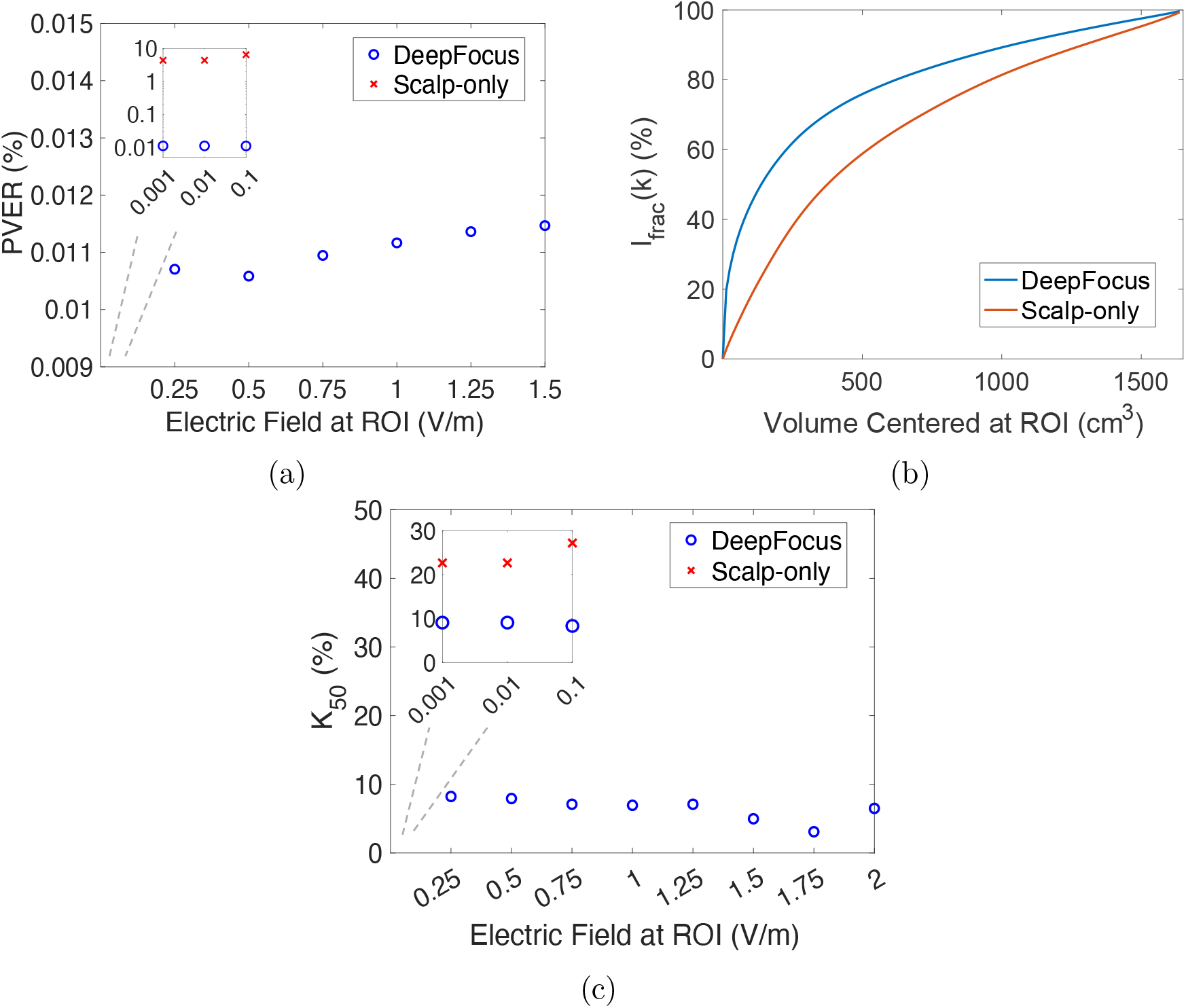
Gain in focality at an OFC target. The current injection patterns were optimized by the Max-Focality Framework. (a) PVER of the field generated by this DeepFocus pattern. (b) Percentage of the total field contained in an increasing volume around the ROI. The desired intensity at the ROI is 0.1V/m. Using DeepFocus, a larger percentage of the total field magnitude is concentrated within a smaller volume centered at the ROI, indicating more focal stimulation at the target. (c) Percentage of voxels around ROI that contain 50% of the total field magnitude (*K*_50_). Note that in (a) and (c), the electric field intensities at the ROI are sampled on a logarithmic scale for smaller values (as scalp-only configurations are unable to attain high fields), and illustrated in the inset plots. Larger intensities are sampled on a linear scale. Both x- and y-axes are on log-scale in the inset in (a).

### 3.2. Brodmann Area 25

BA25 is an important target for drug-resistant depression [2]. As with the OFC, 10 ROIs were randomly selected within BA25 bilaterally. With the maximum-intensity optimization framework in Section 2.4, DeepFocus configurations obtain a maximum of 20.5*×* intensity gain over scalp-only configurations. Fig. 6 and Fig. 7 illustrate a comparison of DeepFocus and scalp-only patterns to target a voxel in BA25. In Fig. 6, an electrode inside the left sphenoid sinus combined with a scalp electrode at Fz are utilized to target the ROI. In scalp-only configurations, due to the medial location of BA25, an anode at Fp1 is used, with a cathode at P4, to create fields traversing through BA25. Relative to the DeepFocus configuration, this optimized scalp-only configuration results in much more diffused fields and undesired activation of superficial brain regions ear the scalp electrodes.

**Figure 6:**
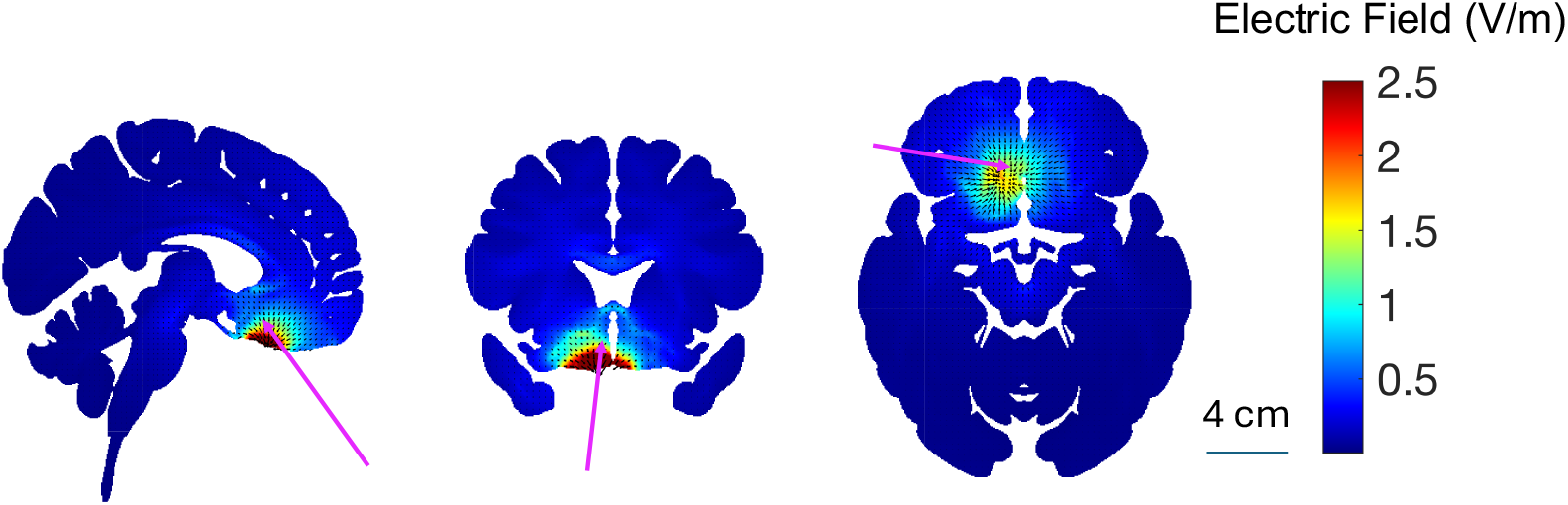
DeepFocus targeting of BA25. Slice views of the electric field magnitude in the brain resulting from 2-electrode stimulation, at the sphenoid sinus and Fz, with 1 mA current. Arrows indicate the location of the ROI.

**Figure 7:**
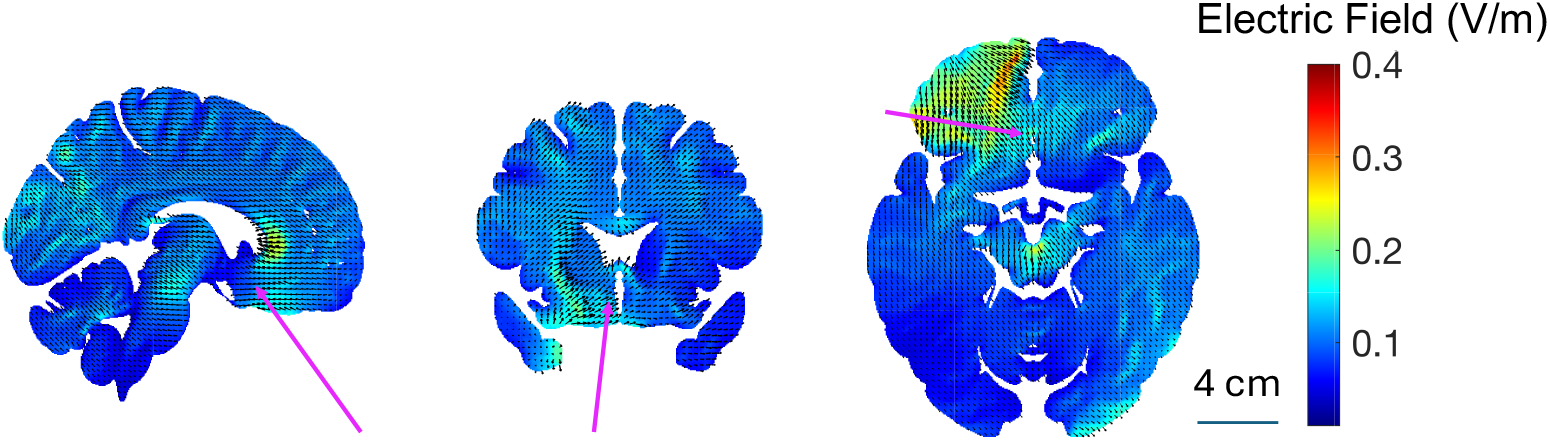
Scalp-only targeting of BA25. Slice views of electric field magnitude in the brain resulting from 2-electrode stimulation (at Fp1 and P4) with 1 mA current.

With the constraint of focality as stated in Section 2.5, we compare the performance of DeepFocus and scalp-only configurations for BA25 targets. As shown in Fig. 8b, 8b, and 8c, the DeepFocus configurations induce more focal fields at BA25 targets.

**Figure 8:**
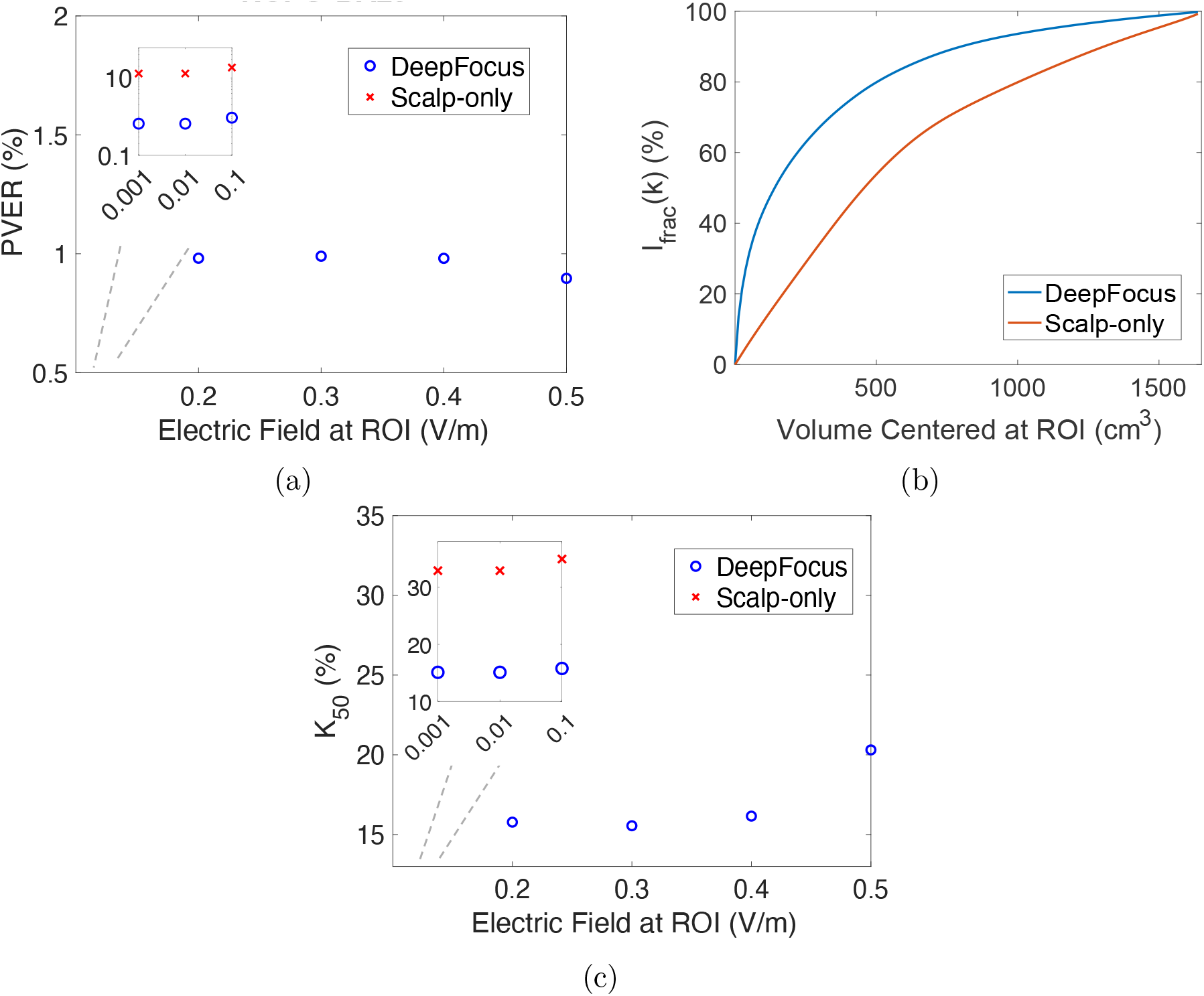
Gain in focality at a BA25 target. The current injection patterns were optimized by the Max-Focality Framework. (a) PVER of the field generated by this DeepFocus pattern.(b)Percentage of total field contained in increasing volumes around the ROI. The desired intensity (*E*_*des*_) at the ROI is 0.1V/m. (c) Percentage of voxels around the ROI that contain 50% of the total field magnitude (*K*_50_).

### 3.3. Summary of Results for Some Reward Circuit Nodes

With the max-focality optimization framework, we searched for the optimal current injection patterns for DeepFocus and scalp-only configurations. Here we report the intensity gain for targeting the ROIs in reward circuit nodes in Table 1, including OFC, BA25, amygdala, nucleus accumbens, anterior ventral thalamus, and anterior hippocampus. 40 ROIs were sampled randomly in the OFC, and 10 ROIs were sampled randomly at other regions shown below. Among the ROIs in OFC, the optimization for both Max-Intensity and Max-Focality tends to select electrodes in the sphenoid sinus for posterior OFC targets, and olfactory cleft electrodes for targets in the anterior OFC (due to their proximity to the OFC). The maximum and median gains are reported.

**Table 1:** The maximum and median gains in intensity at various reward circuit nodes, including the OFC, BA25, amygdala, nucleus accumbens, anterior hippocampus, and anterior ventral thalamus, were assessed. Forty ROIs were sampled within the OFC (i.e., at medial/lateral and anterior/posterior OFC), and 10 ROIs were sampled in each of the other deep brain structures.

| Target ROI | OFC | BA25 | Amygdala | Nucleus Accumbens | Hippocampus | Thalamus |
| --- | --- | --- | --- | --- | --- | --- |
| Maximum Gain | 127.6 | 20.5 | 8.0 | 11.4 | 15.0 | 3.2 |
| Median Gain | 6.7 | 7.7 | 4.3 | 7.9 | 6.6 | 2.3 |

The simulation figures and focality metric plots for each node can be found in the supplementary materials. For each node, we observe a benefit in intensity and focality gain by using DeepFocus configurations, even though the precise benefit is different for different regions based on their proximity to the electrodes.

Further, we include the gain for targeting reward circuit nodes with two additional head models [21, 42], which can be found in the supplementary material Table 1 and Table 2. We post-processed the segmentation of these heads to include the cribriform foramina with the same modifications as those applied to the New York head (i.e., foramina sized 1mm x 1mm). Simulations and optimization were performed under the same conditions (i.e. number of electrodes, electrode size, mesh resolution, regions of interest) for consistency. The results across the three head models are similar. Although the exact gain values, especially the maximum gains, differ due to random sampling of the ROIs, random selection of the desired stimulation direction, and anatomical variations, the median gains for targets remain consistent.

### 3.4. Monte Carlo Simulations

As described in Section 2.7, we modified the cribriform plate anatomy, and across the 10 different head models, we computed the coefficient of variation (i.e., standard deviation*/*mean) of the electric field magnitude for every voxel inside the brain (Fig. 9). Our results suggest that the variation is predominantly limited to the OFC region, where the differences in the foramina sizes and locations induce the most variability. Overall, we observe that the volume of brain voxels that have more than 10% coefficient of variation is 1.4 cm^3^, which is relatively small. If very precise stimulation in medial OFC is desired, this variation could require tailoring to the specific cribriform plate anatomy. In other regions, the effects might be too small to be relevant.

**Figure 9:**
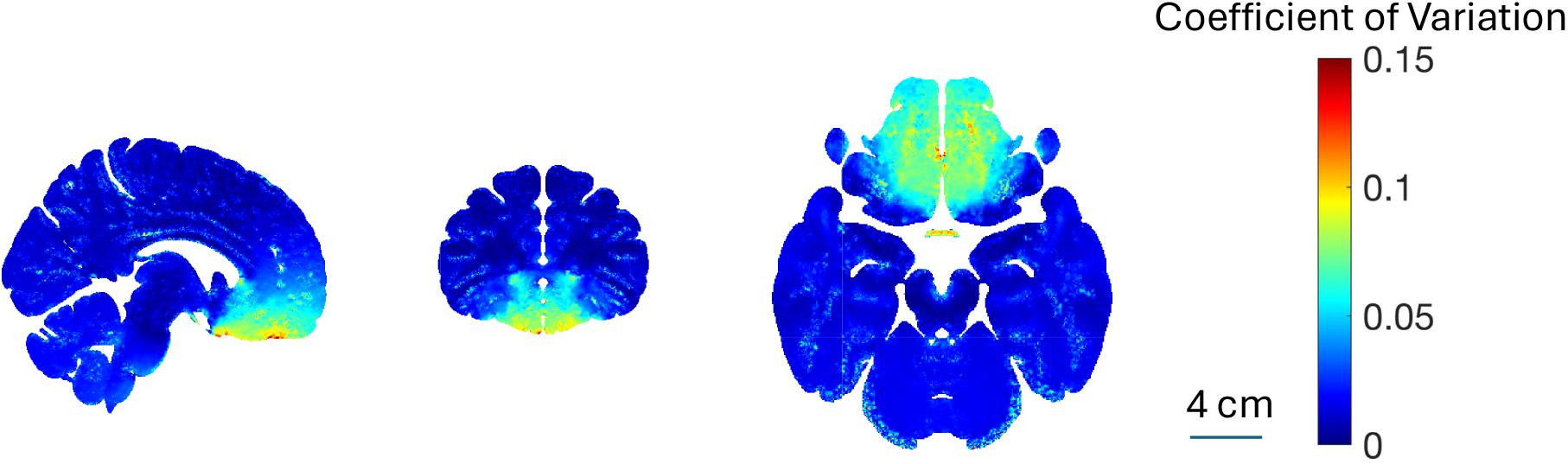
Slice views of coefficient of variation in brain voxels obtained from Monte-Carlo simulation analysis across 10 different head models. The variation of electric field magnitude, due to the variation in locations and numbers of foramina on the cribriform plate, is highly concentrated around the OFC region.

## 4. Cadaver Study Results

The simulation results of Section 3 suggest that DeepFocus patterns create higher intensity and more focal fields than scalp-only configurations. In this section, we report results from cadaver studies that measured the fields achieved in ROIs at the ventral diencephalon and the medial OFC. We quantified the intensity and focality of fields measured in the cadaver experiments and compared them with simulation results. The stimulating and recording electrodes were mapped onto the New York head model through visual inspection of the CT scans. We plotted the *I*_*frac*_ metric (as defined in Section 2.6) to assess focality. Intensity was calculated as the maximum electric field magnitude within the respective volumes. Since conductivity values can vary between in-vivo and ex-vivo conditions [43], we reported the simulated electric field magnitude with the following conductivity values: (i) commonly used ex-vivo values from [21], (ii) ex-vivo conductivity estimates from [44], and (iii) in-vivo estimates at 10 Hz [43], consistent with the frequency of pulses applied in the cadaver study.

### 4.1. Results of Targeting the Ventral Diencephalon

For the ROI at the ventral diencephalon, an anode at the sphenoid sinus and a cathode at Pz were placed based on the Max-Intensity Framework (Fig. 3c). Fig. 10c shows a high field amplitude generated in the target region. The *I*_*frac*_ curves (Fig. 11b) for the simulation and cadaver experiment largely match, indicating similar focality. For 1 mA injected current, the maximum field magnitudes in simulations are 3.9 V/m, 5.2 V/m, and 3.6 V/m corresponding to the three sets of tissue conductivities, consistent with the experimentally measured maximum field magnitude of 4.6 V/m.

### 4.2. Results of Targeting the Medial OFC

For the ROI at the medial OFC, the optimized DeepFocus pattern was an anode inside the olfactory cleft and a cathode at P7, as shown in Fig. 3b. For scalp-only configuration, the optimization pattern was an anode at FP2 and a cathode at P7. Fig. 10 shows the measured electric field magnitude for both (a) DeepFocus and (b) scalp-only patterns for the medial OFC ROI. The DeepFocus pattern created much higher field intensity near the OFC. Fig. 11a shows that the DeepFocus pattern also created more focal fields, in both the cadaver experiment and simulations, at the OFC.

**Figure 10:**
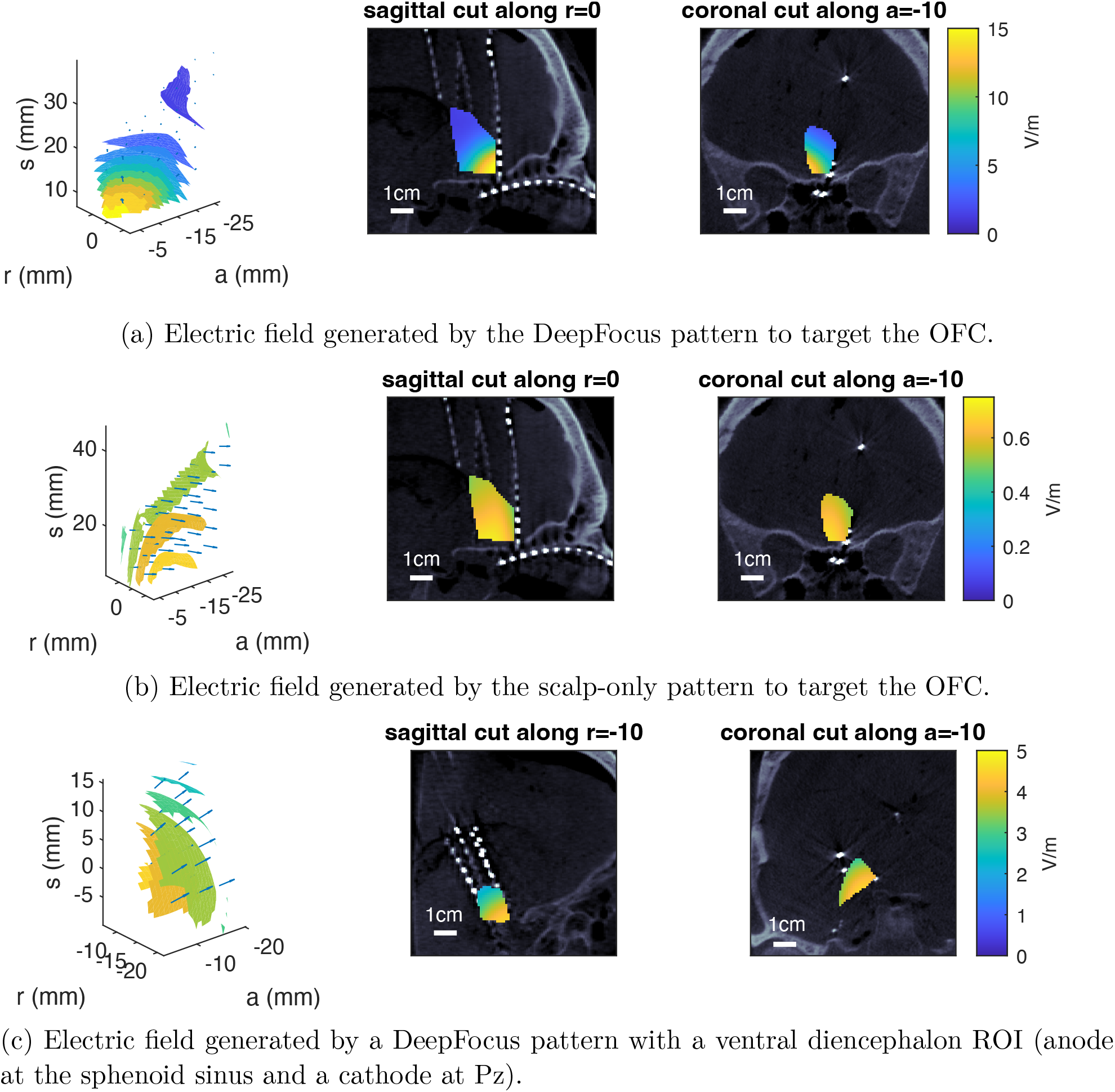
Measured electric field magnitude in cadaveric brain using different electrode configurations for 1 mA injected current. Left: isosurface volume plot; middle: sagittal cut; right: coronal cut (scale bar=1cm). (a): OFC stimulation with the DeepFocus pattern; anode in the olfactory cleft and cathode on the scalp (P7). (b): OFC stimulation with the scalp-only pattern; anode in the scalp (FP2) and cathode on the scalp (P7). (c): Ventral diencephalon stimulation with the DeepFocus pattern; anode in the sphenoid sinus and cathode on the scalp (Pz). Recorded fields are higher intensity and more focal for DeepFocus compared to scalp-only configurations. (a) and (c) use their anode (i.e. the nasal electrode) as the origin; (b) uses the same origin as (a).

**Figure 11:**
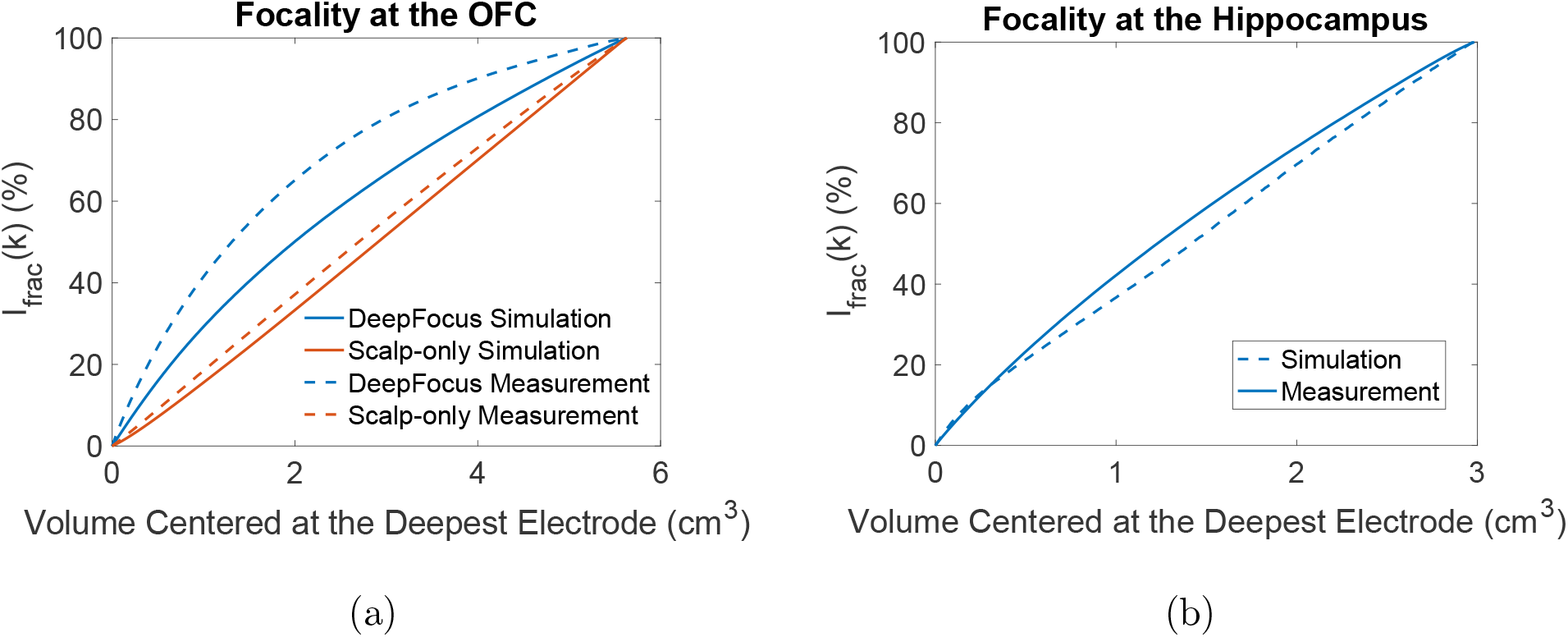
Percentage of total field magnitude constrained in an increasing volume around the deepest recording electrode. (a) Focality of the fields generated by DeepFocus and scalp-only patterns in simulations and experimental measurements. The recorded volume covers the OFC region. (b) Focality of the fields generated by the DeepFocus pattern within the ventral diencephalon region; the *I*_*frac*_ metric for both simulations and experimental measurements was calculated based on the electric field magnitude along the inferior-superior direction.

For 1 mA injected current, the experimentally measured maximum field magnitude was 15.8 V/m, compared to the magnitudes of 4.1 V/m, 5.5 V/m, and 5.7 V/m corresponding to the three sets of conductivity values. Thus, simulations with the three sets of tissue conductivities predict smaller field magnitudes than those measured in the cadaver experiment, although within a factor of 4. This may be due to the cadaver specimen’s cribriform plate being approximately 2.3 times thinner than that of the New York head model, as quantified by examining the cadaver CT scan and the New York head model segmentation.

## 5. Conclusions and Discussion

In this work, we introduced DeepFocus, a minimally-invasive technique to stimulate, e.g., reward circuit nodes, in deep brain regions. We also developed DeepROAST, a platform to simulate fields and optimize electrode locations and currents injected through electrodes for deep-brain stimulation (focal or high-intensity). Through simulations, we compare the efficiency and focality of DeepFocus patterns with traditional scalp-only TES patterns. For the same total injected current, DeepFocus patterns create larger and more focal fields at many different ROIs in the deep-brain. Using cadaver studies, we recorded the electric field induced in the medial OFC by DeepFocus patterns, and observed that it is within a factor of 4 compared to the simulations., and is sufficiently large to create neural response based on estimates from motor cortical stimulation thresholds.

### 5.1. Created field intensities are Likely Sufficient to Create Response at Medial OFC

As noted earlier, medial OFC has been identified as a target for stimulation in major depressive disorder [3]. The medial and lateral OFC play crucial roles in processing reward. While the medial OFC is thought to represent the reward value, the lateral OFC represents non-reward/aversive stimuli [45]. We now discuss how the obtained sEEG measurements, complemented by existing evidence on neural stimulation thresholds, suggest that the fields created are likely sufficient to obtain a neural response at the OFC. Holbrook et al. [29] safely injected 15 mA currents transnasally, with a pulse width of 200-300 µs, establishing safety of these stimulation parameters. With an amplitude of current lower than 15 mA used in Holbrook et al., namely, 10mA used in our cadaver experiment, the medial posterior OFC fields had intensity above 100 V/m.

To assess whether this intensity is sufficient to create neural response, we compare the intensity that suffices in transcranial electrical stimulation using scalp-only electrodes to obtain motor evoked potentials. E.g., in [46], single-pulse TES was conducted for 30 healthy participants, and motor thresholds were estimated. Caulfield et al. also used a 200 µs pulse width, and reported the mean value of the TES motor threshold as 61.35 mA across all participants. Since the relationship between the injected current and the induced electric field is linear, we can use the simulations in [46] to estimate the electric field generated in the primary motor cortex using transcranial stimulation at the average motor threshold. The simulation in Caulfield et al. shows that the electric field at the primary motor cortex reached ∼ 0.5 V/m with 3 mA injected current. With the current at motor threshold (i.e., 61.35 mA), the field intensity at the primary motor cortex would be 10.2 V/m. Our data suggest that the optimized DeepFocus pattern, with 10 mA injected currents, can generate an electric field intensity in the OFC exceeding the 10.2 V/m threshold by more than 10*×*. This makes it plausible that DeepFocus can safely induce a neural response at the OFC, assuming neurons there have a similar excitation threshold to those in the primary motor cortex. This assumption, however, may not be true due to different cytoarchitectures and preferred stimulation directions, which should be informed by future research.

### 5.2. Comparison with Temporal Interference Stimulation

“Temporal Interference” (TI) stimulation was recently proposed as an exciting advance to stimulate the deep brain without stimulating superficial regions [47], and is related to earlier work on interferential stimulation [48, 49]. Rodent studies showed significantly reduced off-target stimulation of deep-brain targets using TI stimulation compared to sinusoidal stimulation, and the technique has received substantial attention [47, 50, 51, 52, 53, 54, 55, 56, 57, 58, 59]. Recent works, however, have revealed several limitations of this technique in being able to perform suprathreshold deep-brain stimulation without superficial effects [51, 60, 61]. In particular, in a recent experimental work [61], our team observed that TI stimulation results in extensive activation of neurons in the superficial brain, and is, fundamentally, a network phenomenon. Surprisingly, cells exhibiting TI stimulation largely fail to continue to exhibit TI stimulation once connections are severed pharmacologically [61]. Related prior computational work from our team [51, 59] as well as others’ [60] show that the required currents for deep-brain TI stimulation are expected to be exceedingly high, likely causing intolerable scalp pain or even tissue damage. This is borne out in experiments: a recent work on human TI stimulation by [62] utilizes currents of 1-2 mA, reporting estimated fields of 0.1 V/m in the deep-brain (for 1 mA injected current), which are insufficient to cause stimulation. The fields created by DeepFocus, as our data suggests, exceed those attained by TI stimulation by 100*×* for the same injected current (*>*10 V/m for 1 mA injected current). The effects produced are thus likely to be very different.

### 5.3. Clinical applications

We envision both chronic and acute applications of DeepFocus. Chronic applications might require persistent stimulation and could be delivered through, e.g. an implant in the sphenoid sinus or ethmoid sinus for stimulating targets of depression treatment. These would require minimally invasive surgery. For acute applications, the treatment could be administered in short sessions with endoscopic insertion and removal of the device. Alternatively, a device could be left in the olfactory cleft or sphenoid sinus for a few days, with sessions conducted by connecting to it as needed. One condition where acute application of DeepFocus could be beneficial in treating substance-use disorder. E.g., circadian rhythms regulate reward processing contributing to substance abuse, and the environment and the time of day play an important role in the de-addiction process [63, 64]. The clinical de-addiction period typically lasts a few weeks, during which substance use is reduced. After the withdrawal period is over and cravings return, treatment is provided alongside environmental management to alter association with cravings [65]. We envision that acute DeepFocus could be applied in this phase, with a clinical practitioner installing the system to stimulate addiction targets, reduce cravings, and disrupt the association of the time of day and/or environmental factors with cravings.

### 5.4. Anatomical Complexity and Individual Variations in the Skull-Base

In the DeepROAST platform, we manually modified the tissue mask to include anatomical details on the cribriform plate. However, the areas, locations and sizes of holes on the cribriform plate vary across individuals and ages. In Section 3.4, we conducted Monte Carlo simulations and observed that the change of field magnitude due to the variations in cribriform foramina anatomy appears to be substantial only in the OFC region, and yet, it is well within an order of magnitude. However, the Monte Carlo simulations may also be insufficient due to our assumptions on possible variations. Future work will investigate data-driven approaches for forward matrix estimation/correction regardless of the cribriform plate anatomy.

### 5.5. Safety of Transnasal Stimulation

Although accessing the sphenoid sinus transnasally is a common surgical procedure, e.g. for pituitary tumor removal [66], the safety and tolerability of current amplitudes and waveforms that can be injected from the sphenoid sinus are unknown, and should be informed by future animal and human studies. In practice, this may require slowly ramping up the current, and monitoring physiological parameters to ensure safety (as is commonly done in experimental DBS procedures [2]). Similar to the olfactory cleft, the sphenoid sinus is also lined with mucosa, which may contribute to site-dependent variation in impedance [33]. In addition, since the sphenoid sinus is physically proximal to the optic chiasma, stimulation could create visual artifacts, which may limit some applications, but could also help calibrate the current amplitude and the pattern choice.

### 5.6. Discomfort due to Transnasal Stimulation

This work is based on simulation and cadaver studies, where the discomfort of electrode placement and stimulation cannot be assessed. However, we can be informed by common practices in endonasal procedures, as well as previous studies that attempted transnasal stimulation. Commonly, local anesthetics, such as tetracaine, are used to avoid discomfort in endonasal procedures. Straschill et al. [30] and Ishimaru et al. [31] used nasal spray (e.g. 0.1% epinephrine) for vasoconstriction, which allowed easier access of electrodes to the olfactory cleft. In Holbrook et al. [29], with 3 mm electrode diameter, a maximum of 15 mA was safely injected (pulse width of 0.2-0.3 msec) in the olfactory cleft. In Ishimaru et al. [31], smaller currents were injected (up to 2mA), but longer pulse duration was used (up to 0.5 msec). The placement and stimulation in the olfactory mucosa only produced a tactile-like somatic sensation but not pain in all five participants. Even these somatic sensations could be easily avoided by moving the electrode position. Placement of electrodes in the respiratory mucosa caused pain, but not in the olfactory mucosa, where we propose to stimulate. Further, Weiss et al. [33] also found differences in sensations across individuals, sites, and stimulation frequencies inside the olfactory cleft. They also note that transnasal electrodes are in contact with the physiologically varying mucosa, leading to fluctuating impedance, which, in practice, may result in current or voltage spikes. These factors will be important as DeepFocus approaches clinical practice.

### 5.7. Choice of Optimal Orientation of Stimulation

For the regions of interest considered in this paper, the optimal orientations of stimulating fields are assumed to be random. Optimal field directions are sophisticated functions of surrounding circuitry, and improving on these assumptions requires further studies that investigate optimal field orientations for neurons in different reward circuit nodes, e.g. in animal models, using non-invasive-like stimulation (i.e., stimulation with fields that have low gradients). This can be easily specified in the optimization framework as informed by future research.

### 5.8. Cadaver Study Limitations

In our cadaver study, only one specimen was investigated. Due to the variations in the skull base anatomy, it is important to increase the sample size so that the intensity and focality gain of DeepFocus can be quantified statistically. It is also known that, while within an order of magnitude, conductivity of cadaver specimens can differ from live tissue [67, 68]. In Section 4, we compared our simulation results with field data recorded in cadavers. Although simulated electric field intensity is within an order of magnitude with the recording, there is ∼3x difference for olfactory cleft stimulation. This may be due to several contributing factors: (i) variation of tissue conductivities, as described in Section 4; (ii) the specimen in our experiment was fresh-frozen and thawed, which may contribute to changes in conductivities. (iii) the cribriform plate of the cadaver specimen is thinner than the head model we used in DeepROAST, which may contribute to a larger measured electric field at the OFC than predicted by simulations.

## Acknowledgments

We thank Dr. Eric Wang, an otolaryngologist and an expert in endonasal and skull-base procedures, for helpful discussions on the feasibility of installation of olfactory cleft and sphenoid sinus electrodes, and Dr. Alex Yu, a skull-base neurosurgeon at AHN, on potential applications of this approach in functional neurosurgery. We thank Dr. Raghav Shah and Dr. Khaled Moussawi, clinicians and de-addiction experts, on discussions regarding potential use in treatment of substance-use disorder. We also thank Jenn Gooch for help with Figure 1. This work was supported by The Assistant Secretary of Defense for Health Affairs endorsed by the Department of Defense, in the amount of ($258,606), through the Peer Reviewed Medical Research Program under Award Number HT9425-24-1-0018. Opinions, interpretations, conclusions, and recommendations are those of the author(s) and are not necessarily endorsed by The Assistant Secretary of Defense for Health Affairs endorsed by the Department of Defense. YG is supported by the Centre for Machine Learning and Health (CMLH) fellowship at Carnegie Mellon University. This material is also based upon work supported by the Chuck Noll Foundation for Brain Injury Research, and the Naval Information Warfare Center (NIWC) Atlantic and the Defense Advanced Research Projects Agency (DARPA) under Contract No. N65236-19-C-8017. Any opinions, findings and conclusions or recommendations expressed in this material are those of the author(s) and do not necessarily reflect the views of the NIWC Atlantic and DARPA.

