## Supporting Information for "DeepFocus: A Transnasal Approach for Optimized Deep Brain Stimulation of Reward Circuit Nodes"

### Supplementary Materials

#### 1 Results for Reward Circuit Targets

In this section, we provide simulation results for the electric field generated by DeepFocus patterns with the Max-Intensity Framework. We also include results withthe Max-Focality Framework, emphasizing on the focality metric.

Overall, we observe that, with Max-Intensity optimization, DeepFocus patterns are able to attain larger fields for deep brain targets, namely, the amygdala, nucleus accumbens, anterior ventral thalamus, and anterior hippocampus. Slice views of the generated fields at these targets are provided in Fig. 1, 3, 5, and 7. The results of Max-Focality optimization at these targetes are provided in Fig. 2, 4, 6, and 8. All the results for reward circuit nodes are reported with a nominal total injected current of 1mA. As discussed in the main manuscript, currents larger than 1mA have been injected safely in the olfactory cleft in prior work. To obtain fields for any current, the electric field can be obtained easily as it scales linearly with the injected current.

Fig. 11 and Fig. 12 provide slice views of Max-focality optimized electric field with DeepFocus and scalp-only patterns to target an ROI at the orbitofrontal cortex. In general, the Max-Focality Framework utilizes more electrodes than the Max-Intensity Framework. When both are optimized with the Max-Focality Framework, scalp-only patterns generate more diffused fields compared with DeepFocus patterns. Fig. 13 illustrates the slice views of field generated near the active electrode, where current injection pattern is identical to that of Fig. 4c in the paper.

To test sensitivity to specific head models, we conducted simulation and optimization on two more head models from [1, 2], and summarized the gains in reward circuit nodes in Table 1 and Table 2. The focality figures for targeting an orbitofrontal cortex ROI are included in Fig. 9 and Fig. 10. DeepFocus continues to create larger and more focal fields in these two head models, consistent with the observation with the New York head model.

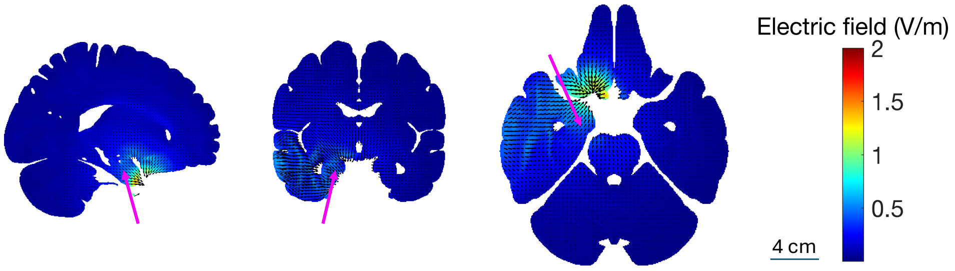


Figure 1: Slice views of the electric field created by the DeepFocus Max-Intensity pattern (anode: sphenoid sinus, cathode: T7) to target an amygdala ROI. The resulted field intensity is above 0.6 V/m at amygdala along the desired direction, and is focal around the ROI even without explicit focality constraints.

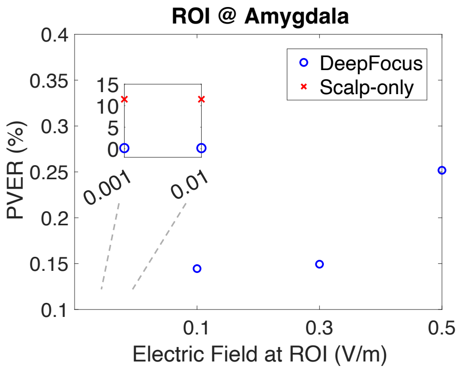


(a)

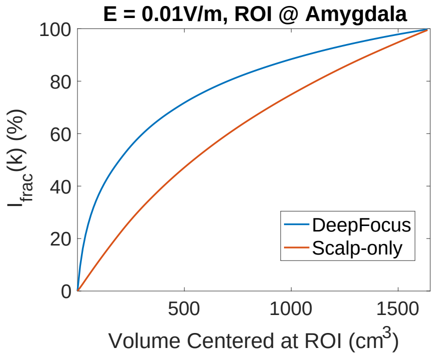


(b)

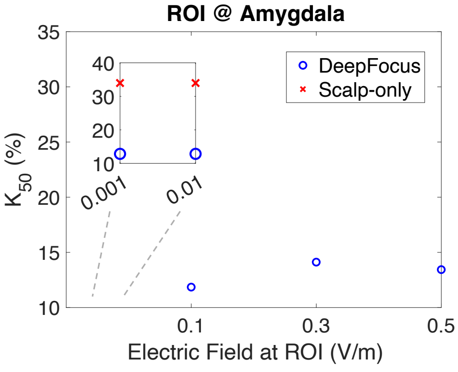


(c)

Figure 2: Gain in focality at an amygdala target with the Max-Focality optimization framework. (a) PVER of the field generated by this DeepFocus pattern. (b) Percentage of total field contained in an increasing volume around the ROI. (c) Percentage of voxels around the ROI that contain 50% of the total field magnitude.

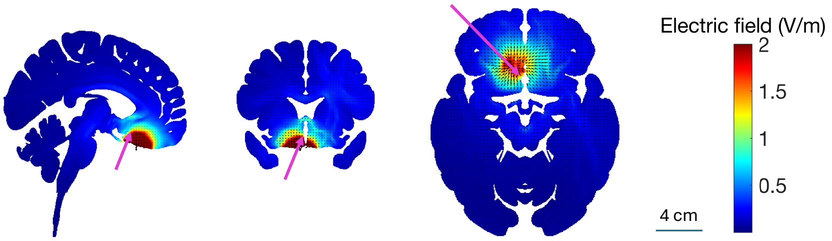


Figure 3: Slice views of the electric field created by the DeepFocus Max-Intensity pattern (anode: left sphenoid sinus, cathode: FC4) to target a nucleus accumbens ROI. The electric field reaches 1.4 V/m at the ROI with 1mA total injected current.

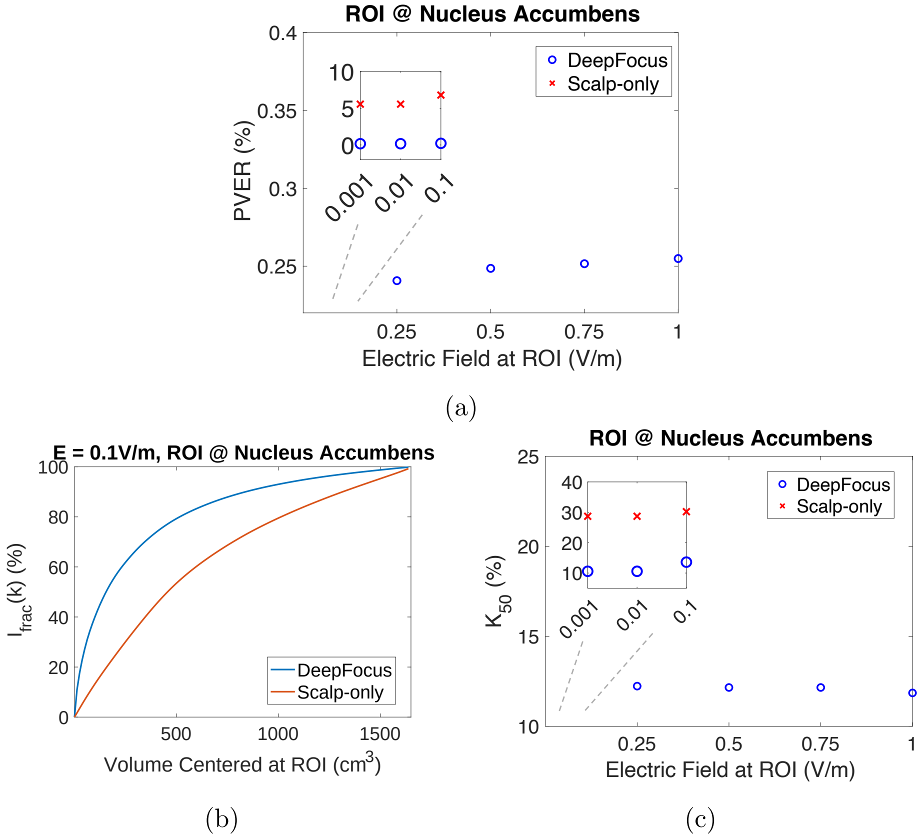


Figure 4: Gain in focality at a nucleus accumbens target with the Max-Focality optimization framework. (a) PVER of the field generated by this DeepFocus pattern. (b) Percentage of total field contained in an increasing volume around the ROI. (c) Percentage of voxels around the ROI that contain 50% of the total field magnitude.

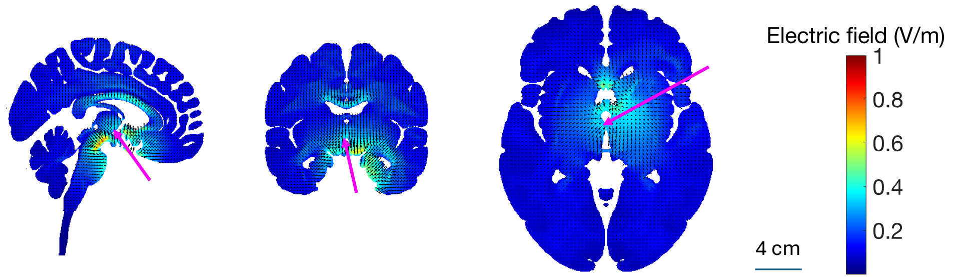


Figure 5: Slice views of the electric field created by the DeepFocus Max-Intensity pattern (anode: sphenoid sinus, cathode: Cz) to target an anterior ventral thalamus ROI. The electric field intensity reaches 0.29 V/m at the ROI with 1mA total injected current, which is smaller than that of other regions since thalamus is physically far away from the transnasal electrodes.

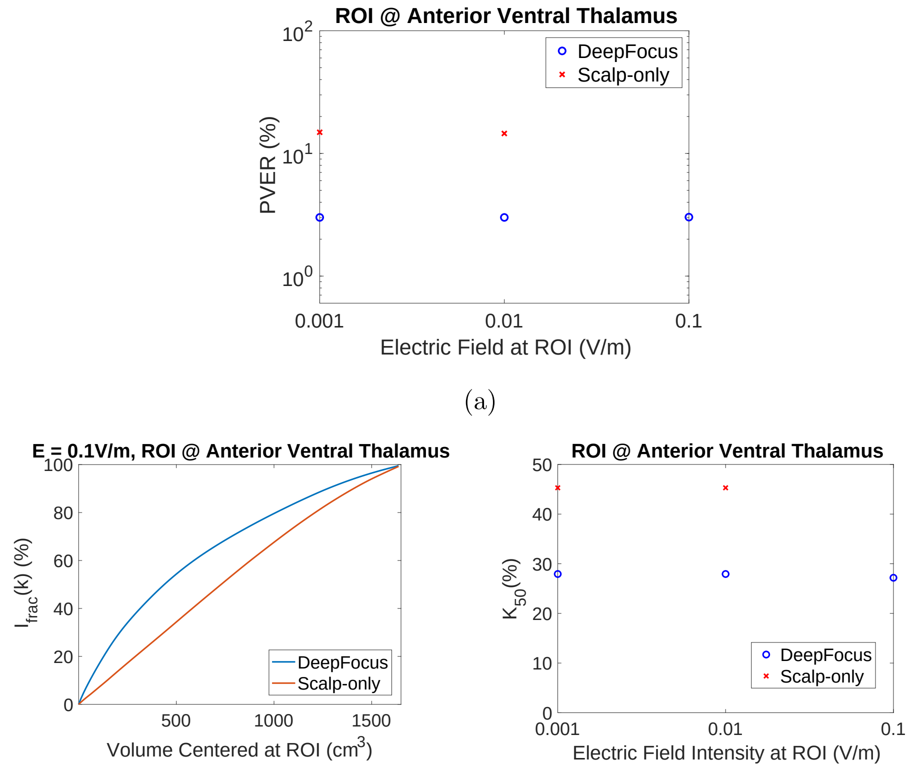


(a)

(b) (c)

Figure 6: Gain in focality at an anterior ventral thalamus target with the Max-Focality optimization framework. (a) PVER of the field generated by this DeepFocus pattern. (b) Percentage of total field contained in an increasing volume around the ROI. (c) Percentage of voxels around the ROI that contain 50% of the total field magnitude.

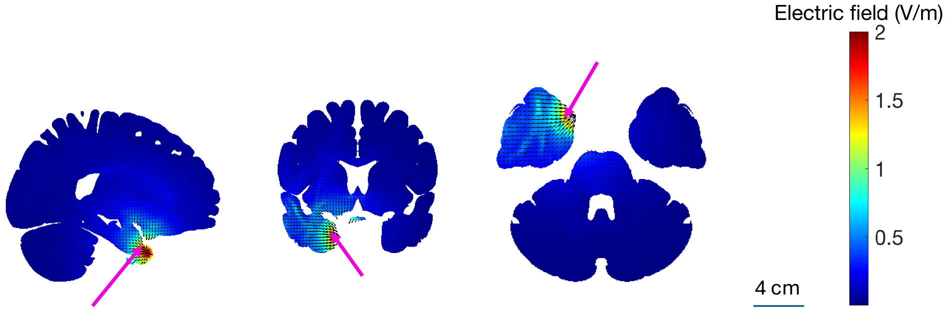


Figure 7: Slice views of the electric field created by the Max-Intensity Deep-Focus pattern (anode: sphenoid sinus, cathode: T7) to target an anterior hippocampus ROI. The electric field reaches 1.2 V/m at the ROI along the desired direction with 1mA total injected current.

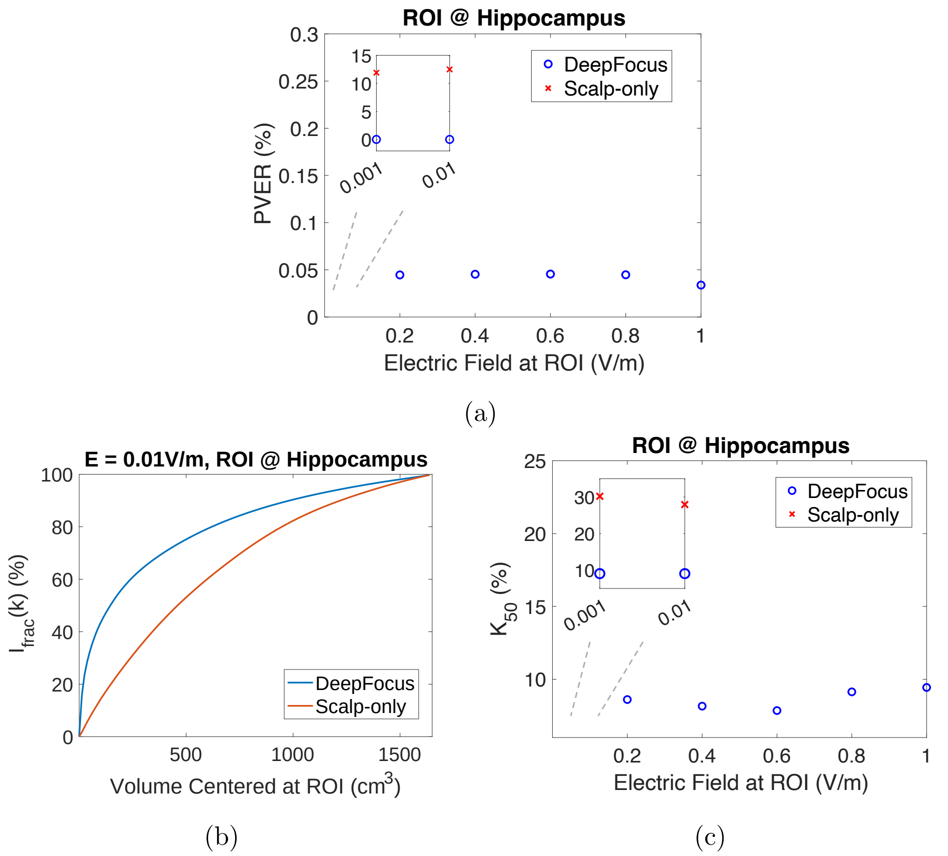


Figure 8: Gain in focality at a hippocampus target with the Max-Focality optimization framework. (a) PVER of the field generated by this DeepFocus pattern.(b) Percentage of total field contained in an increasing volume around the ROI. (c) Percentage of voxels around the ROI that contain 50% of the total field magnitude.

| Target ROI | OFC | BA25 | Amygdala | Nucleus Accumbens | Hippocampus | Thalamus |
|---|---|---|---|---|---|---|
| Maximum Gain | 87.6 | 13.8 | 6.6 | 12.8 | 10.7 | 5.2 |
| Median Gain | 6.9 | 6.2 | 4.2 | 7.0 | 5.9 | 3.5 |

Table 1: Intensity gain in some reward circuit nodes with the MIDA head model.

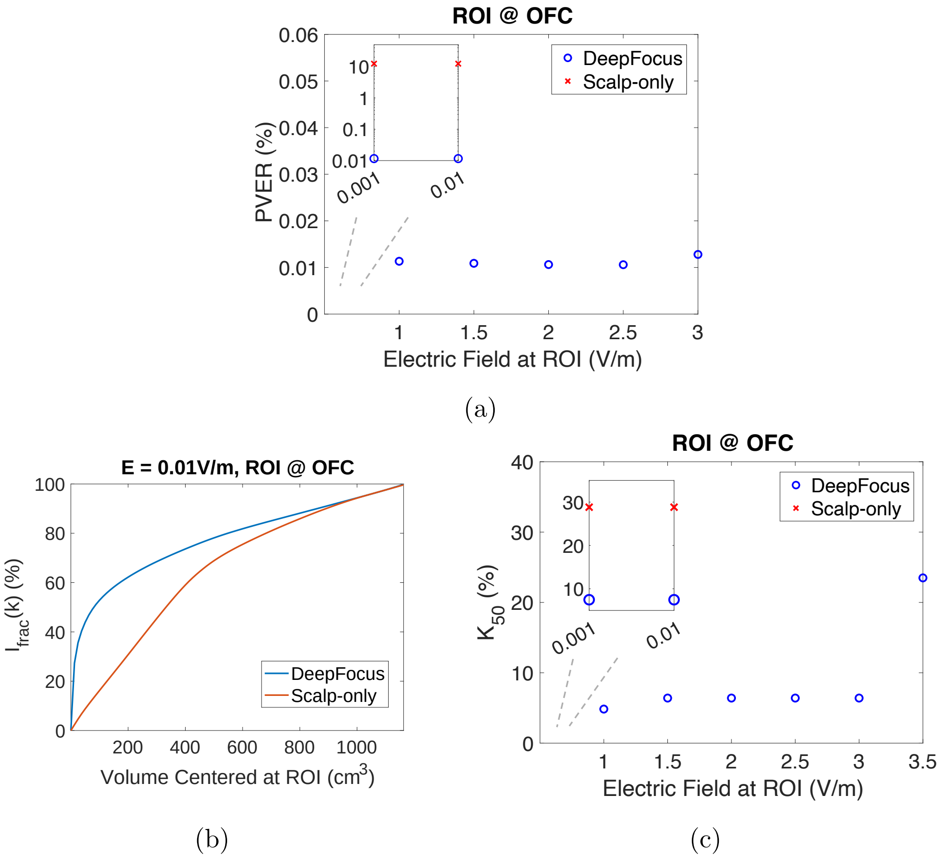


Figure 9: Gain in focality at an OFC target ROI with the MIDA head model utilizing the Max-Focality optimization framework. (a) PVER of the field generated by this DeepFocus pattern. (b) Percentage of total field contained in an increasing volume around the ROI. (c) Percentage of voxels around the ROI that contain 50% of the total field magnitude.

| Target ROI | OFC | BA25 | Amygdala | Nucleus Accumbens | Hippocampus | Thalamus |
|---|---|---|---|---|---|---|
| Maximum Gain | 93.2 | 20.8 | 7.9 | 7.5 | 9.0 | 4.3 |
| Median Gain | 5.6 | 5.4 | 4.9 | 4.4 | 6.1 | 2.9 |

Table 2: Intensity gain in some reward circuit nodes with the Subject 1 head model.

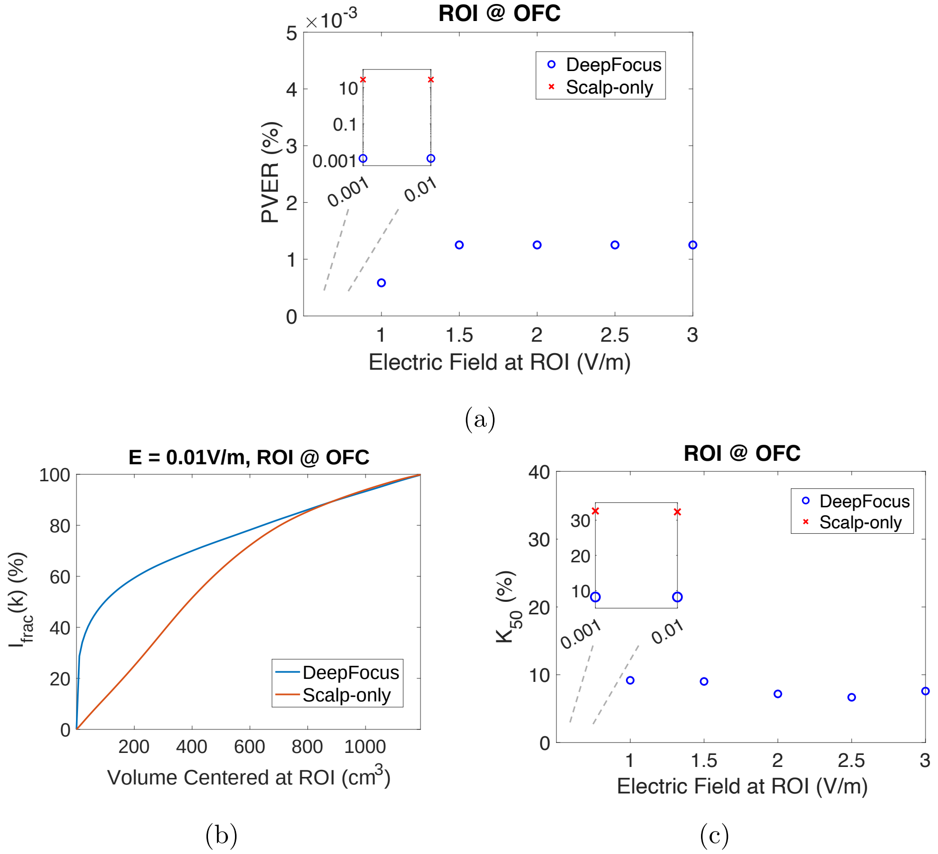


Figure 10: Gain in focality at an OFC ROI with the Subject1 head model utilizing Max-Focality optimization framework. (a) PVER of the field generated by this DeepFocus pattern. (b) Percentage of total field contained in an increasing volume around the ROI. (c) Percentage of voxels around the ROI that contain 50% of the total field magnitude.

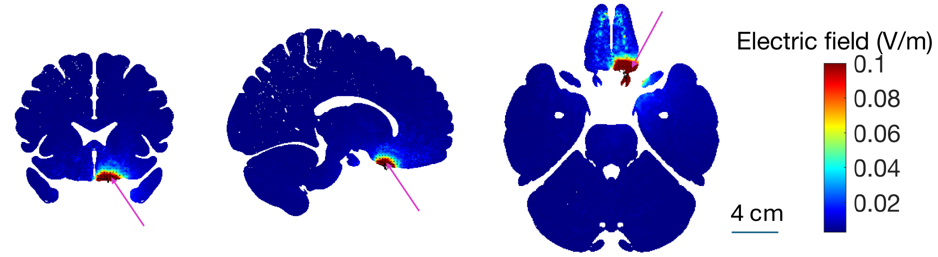


Figure 11: Slice views of the electric field generated by DeepFocus. The current injection pattern was optimized by the Max-focality Framework to target the OFC. Three electrodes in the olfactory cleft in both hemispheres (six in total) are utilized, along with 2 electrodes in the right sphenoid sinus. The electric field intensity along the desired direction at the ROI is 0.1 V/m.

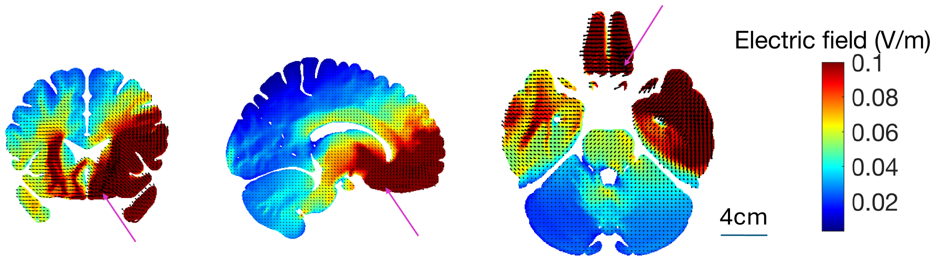


Figure 12: Slice views of the electric field generated by scalp-only electrodes. The current injection pattern was optimized by the Max-focality Framework to target the OFC. The electrodes with the largest current injections are FT8, TP8, F7, F4, FC4, and FP1. The electric field intensity along the desired direction at the ROI is 0.1 V/m.

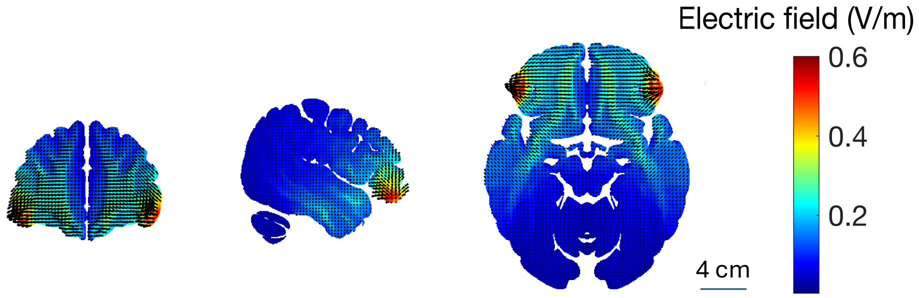


Figure 13: Slice views of the electric field generated by the Max-Intensity scalp-only electrodes (F7 and F8) to target the OFC. The slices are shown close to the anode instead of the ROI. This optimized scalp-only pattern stimulates the shallow brain regions more than deep brain targets.

#### 2 Simulations with Different Conductivities

Fig. 14 and Fig. 15 present simulation results for current injection patterns used in our cadaver experiments, with different sets of conductivity values. The total injected current is 1mA.

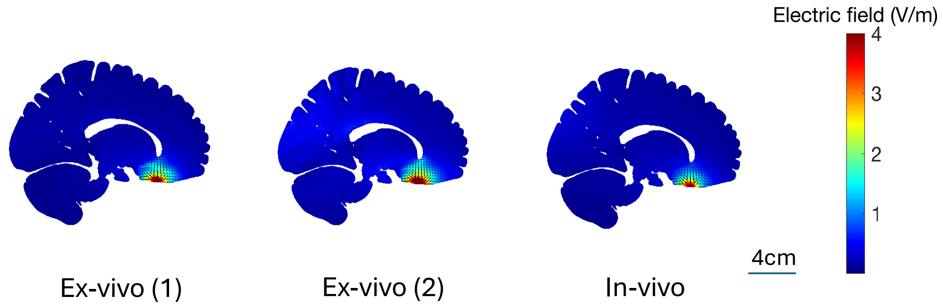


Figure 14: Simulated electric field generated by 3 sets of conductivity values with the cathode at sphenoid sinus and the anode at Pz. Ex-vivo (1) conductivities are from [2], Ex-vivo (2) conductivities are from [3], and in-vivo conductivities are from [4]. The total injected current is 1mA.

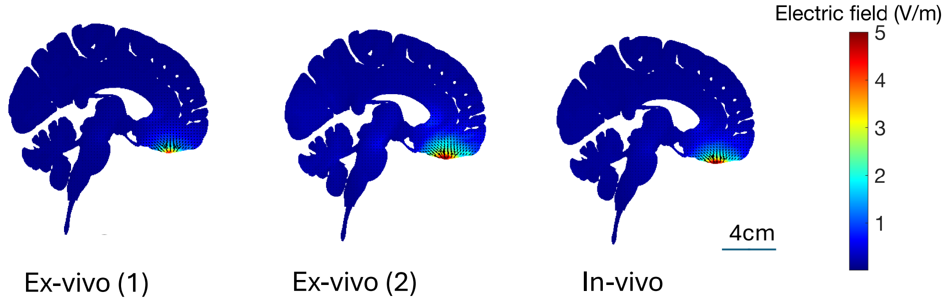


Figure 15: Simulated electric field generated by 3 sets of conductivity values with cathode at olfactory cleft and anode at P7. Ex-vivo (1) conductivities are from [2], Ex-vivo (2) conductivities are from [3], and in-vivo conductivities are from [4]. The total injected current is 1mA.

#### 3 Visualization of the Monte Carlo Samples

Fig. 16 shows the horizontal views of the nasal region in the Monte Carlo simulation samples. The number and locations of the foramina were randomly sampled.

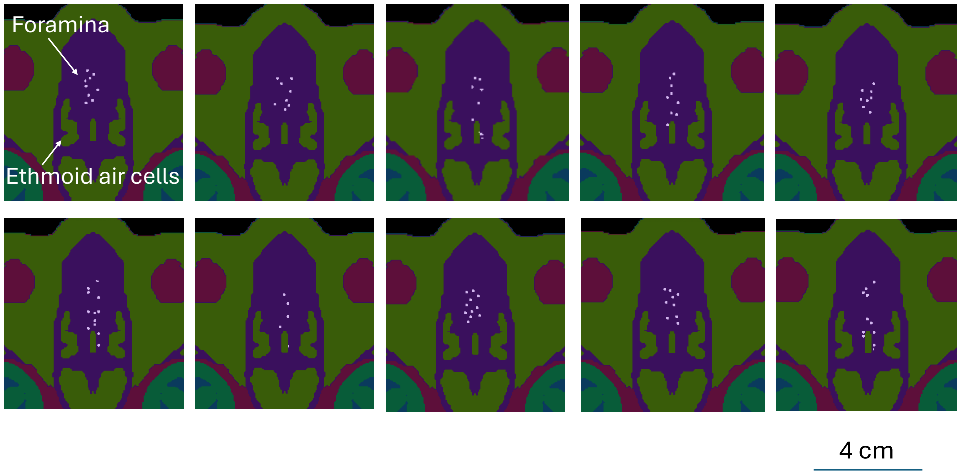


Figure 16: The 10 randomly generated Monte Carlo simulation samples of foramina on the cribriform plate that were utilized in our Monte Carlo analysis. The cribriform plate was segmented, and a random number of foramina were placed at random locations on the cribriform plate.

#### References

[1] Maria Ida Iacono, Esra Neufeld, Esther Akinnagbe, Kelsey L. Bower, Johanna Wolf, Ioannis Vogiatzis Oikonomidis, Deepika Sharma, Bryn A. Lloyd, Bertram Jakob Wilm, Michael Wyss, Klaas Paul Pruessmann, András Jakab, Nikos Makris, Ethan D Cohen, Niels Kuster, Wolfgang Kainz, and Leonardo M. Angelone. Mida: A multimodal imaging-based detailed anatomical model of the human head and neck. *PLoS ONE*, 10, 2015.

[2] Yu Huang, Abhishek Datta, Marom Bikson, and Lucas C. Parra. Realistic volumetric-approach to simulate transcranial electric stimulation—ROAST—a fully automated open-source pipeline. *Journal of neural engineering*, 16:056006 – 056006, 2017.

[3] C. Gabriel, S. Gabriel, and E. Corthout. Compilation of the dielectric properties of body tissues at rf and microwave frequencies. *Physics in Medicine and Biology*, 41(11):2231–2249, 1996.

[4] Timothy Andrew Wagner, Uri T. Eden, Jarrett Rushmore, Christopher J. Russo, Laura Dipietro, Felipe Fregni, Stephen Simon, Stephen Rotman, Naomi B. Pitskel, Ciro Ramos-Estebanez, Álvaro Pascual-Leone, Alan J. Grodzinsky, Markus Zahn, and Antoni Valero-Cabré. Impact of brain tissue filtering on neurostimulation fields: A modeling study. *NeuroImage*, 85:1048–1057, 2013.
